# Basal ppGpp signalling by SpoT integrates metabolism with acid resistance

**DOI:** 10.64898/2026.01.26.700336

**Authors:** Yueyue Liu, Muriel Leandra Schicketanz, Xichuan Zhai, Ling Deng, Kenn Gerdes, Yong Everett Zhang

**Author notes:** Correspondence: Yong Everett Zhang. Y.Y.L. and M.L.S. contributed equally to this work.

## Abstract

Bacteria maintain low basal levels of the alarmone ppGpp during steady-state growth, yet how this basal state is established and why it matters physiologically remain poorly understood. Here, we show that basal ppGpp represents an actively maintained regulatory state that coordinates metabolic homeostasis with stress resistance. Using a targeted perturbation of SpoT regulation in *Escherichia coli*, we uncover a sharp requirement for a sub-basal yet non-zero ppGpp pool to sustain growth in minimal medium and survival under extreme acid stress. Disruption of this basal state leads to misallocation of metabolic flux into arginine biosynthesis, depletion of glutamate, and collapse of the glutamate-dependent Gad acid resistance system. Through intragenic suppressor analysis, enzymatic perturbations, and protein-level feedback measurements, we further demonstrate that SpoT intrinsically tunes basal ppGpp through a distributed intramolecular regulatory network coupled to negative feedback control of SpoT abundance. This regulatory logic stabilizes ppGpp within a narrow physiological window, below the threshold of canonical stringent response activation. The requirement for SpoT-dependent basal ppGpp regulation is conserved in pathogenic *Salmonella* and *Shigella*. Together, our findings establish basal ppGpp as a distinct and actively regulated signaling regime that integrates metabolism and stress preparedness beyond acute starvation responses.

## Introduction

Bacteria inhabit environments that fluctuate rapidly and often impose multiple, simultaneous stresses. Successful survival under such conditions requires tight coordination between core metabolic processes and stress-protective responses in a manner that preserves essential resources while maintaining viability. For enteric bacteria in particular, the ability to balance nutrient availability with resistance to extreme acidity - such as that encountered during gastric passage - is a key determinant of ecological fitness and pathogenic potential^1,2^. Despite extensive study of individual stress-response pathways, how bacteria integrate metabolic homeostasis with acid resistance remains incompletely understood.

A central regulator of bacterial metabolic adaptation is the stringent response, a conserved signaling network mediated by the alarmone guanosine tetra- and pentaphosphate (ppGpp)^3–6^. Canonically, ppGpp is produced in large bursts upon amino acid starvation, when uncharged tRNAs activate the ribosome-associated enzyme RelA in *Escherichia coli*^7–11^. This acute increase in ppGpp rapidly remodels transcription, repressing stable RNA synthesis while redirecting metabolic flux toward amino acid biosynthesis^12–16^. In parallel, ppGpp directly targets numerous metabolic enzymes, rebalancing nucleotide and amino acid pools^17–21^. This RelA-driven stringent response has therefore been widely viewed as an emergency program that facilitates rapid bacterial adaptation to acute nutrient stress.

Once this acute response has restored metabolic balance and growth resumes, ppGpp levels decline but do not return to zero. In contrast to the well-characterized high-amplitude stringent response, much less is known about the physiological role of these low, basal levels of ppGpp that persist during adapted growth^22^. Basal ppGpp is believed to be primarily controlled by SpoT, a bifunctional enzyme that both synthesizes and hydrolyzes ppGpp^23^ and responds to diverse environmental cues distinct from amino acid starvation^24–30^. Genetic and physiological studies have established that SpoT activity is essential for viability, virulence, and stress adaptation across many bacterial species^3,31,32^. However, because SpoT’s hydrolase activity is required to counterbalance RelA-mediated ppGpp synthesis, direct analysis of SpoT regulatory function has been challenging. As a result, basal ppGpp has often been treated as a passive background signal rather than an active regulatory regime with distinct physiological roles.

Acid resistance provides a compelling context in which to examine basal ppGpp regulation. *E. coli* is capable of surviving exposure to pH values as low as 2.5 through multiple acid resistance systems, among which the glutamate-dependent Gad system is the most effective^2,33,34^. This system uses cytoplasmic glutamate decarboxylases (GadA and GadB) to convert glutamate to γ-aminobutyrate, consuming intracellular protons, while the GadC antiporter exchanges γ-aminobutyrate for extracellular glutamate to sustain the cycle^35,36^. The Gad system is subject to complex regulation involving transcriptional activators (GadE), global stress regulators (e.g., ppGpp), and growth-phase signals (e.g., RpoS)^2,36^. Notably, glutamate itself occupies a central position in cellular metabolism, serving as a key nitrogen donor and metabolic hub. How bacteria coordinate glutamate homeostasis with glutamate-dependent acid resistance under nutrient-limited conditions remains poorly understood.

Here, we set out to investigate how SpoT-dependent basal ppGpp signaling contributes to the integration of amino acid metabolism with acid resistance during nutrient-limited growth. As an entry point into this poorly understood regulatory regime, we focused on a strictly conserved histidine residue (H414) within the C-terminal regulatory region of SpoT. Perturbation of this residue resulted in pronounced physiological imbalances, including altered metabolic flux through arginine biosynthesis, depletion of intracellular glutamate, and reduced expression of the glutamate-dependent Gad acid resistance system. These observations provided a framework for systematically dissecting how SpoT-dependent basal ppGpp supports metabolic and stress-response homeostasis.

Because SpoT homologs are conserved across enteric bacteria, we further asked whether the regulatory principles revealed by this approach extend beyond *E. coli*. Addressing this question allowed us to evaluate whether SpoT-dependent basal ppGpp represents a species-specific phenomenon or a more general regulatory regime. In this context, our study seeks to expand the functional scope of ppGpp signaling beyond RelA-mediated acute starvation responses to include a role in coordinating metabolism and stress resistance during nutrient-limited growth.

## Results

### Perturbation of SpoT regulation leads to severe growth defects in minimal medium

To examine how SpoT-dependent regulation supports growth under nutrient-limited conditions independently of RelA, we focused on the conserved C-terminal regulatory region of SpoT, which is highly conserved among RelA/SpoT homologs but remains poorly understood mechanistically. Sequence alignment identified a strictly conserved histidine residue (H414 in *E. coli* SpoT) within the TGS domain, corresponding to H432 in RelA (**Extended Data Fig. 1a, 1b**), a residue cucial for sensing amino acid starvation^8–11^.

As a genetic probe of this conserved regulatory region, we generated a SpoT H414A substitution in a *ΔrelA* background. While the *ΔrelA spoT^wt^* (hereafter referred to as *spoT^wt^*) strain grew normally, the H414A mutant (*spoT^H414A^*) displayed a severe and unexpected growth defect on glucose minimal medium (M9Glc), both on solid and in liquid culture (**Fig. 1a, 1b**). This unanticipated phenotype indicated that SpoT H414 plays a critical role in supporting growth under nutrient-limited conditions and prompted further investigation into its underlying molecular basis.

**Figure 1.**
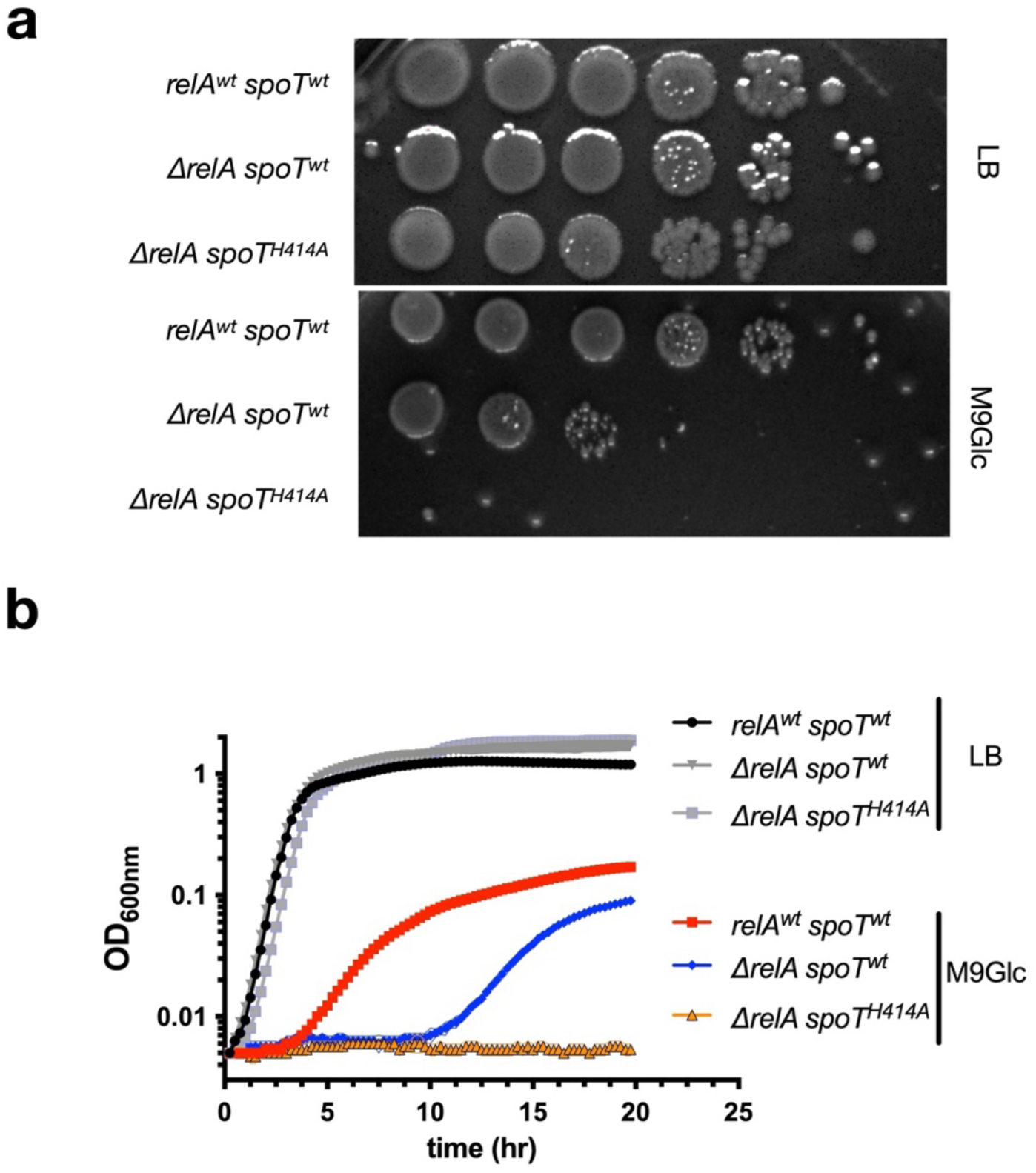
A conserved SpoT H414A mutation reveals defective growth on M9Glc medium. **(a)** Growth of wild type *E. coli* MG1655 (*relA^wt^ spoT^wt^*), and *ΔrelA spoT^wt^*, *ΔrelA spoT^H414A^* strains on LB and M9Glc solid media. **(b)** Growth dynamics of the same strains in liquid LB and M9Glc media.

### Loss of SpoT regulation disrupts transcriptional programs for stress and acid resistance

To gain insight into the molecular defects underlying the growth failure of *spoT^H414A^* cells, we performed transcriptomic analysis of the *spoT^H414A^* and *spoT^wt^* strains. Because *spoT^H414A^* cells failed to proliferate in M9Glc, cultures were incubated under identical conditions until the *spoT^wt^* strain entered early exponential phase, at which point RNA was harvested from both strains (**Extended Data Fig. 1c**).

This analysis revealed extensive transcriptional dysregulation in *spoT^H414A^* cells, with strong repression of stress adaptation and acid resistance genes. Among the most prominently downregulated loci were the glutamate-dependent Gad acid resistance system (*gadE*, *gadX*, *gadW*, *gadA*, *gadB*, *gadC*), as well as additional components of the acid fitness island, including the periplasmic chaperones *hdeAB* (**Fig. 2a, 2b**). Genes associated with osmotic and general stress responses were similarly reduced; in contrast, genes linked to phage shock and SOS responses were strongly induced (**Extended Data Fig. 1d-f**), indicating a broad physiological imbalance. The pronounced repression of acid resistance pathways indicated a defect in acid survival and prompted direct physiological testing of *spoT^H414A^* cells under acidic conditions.

**Figure 2.**
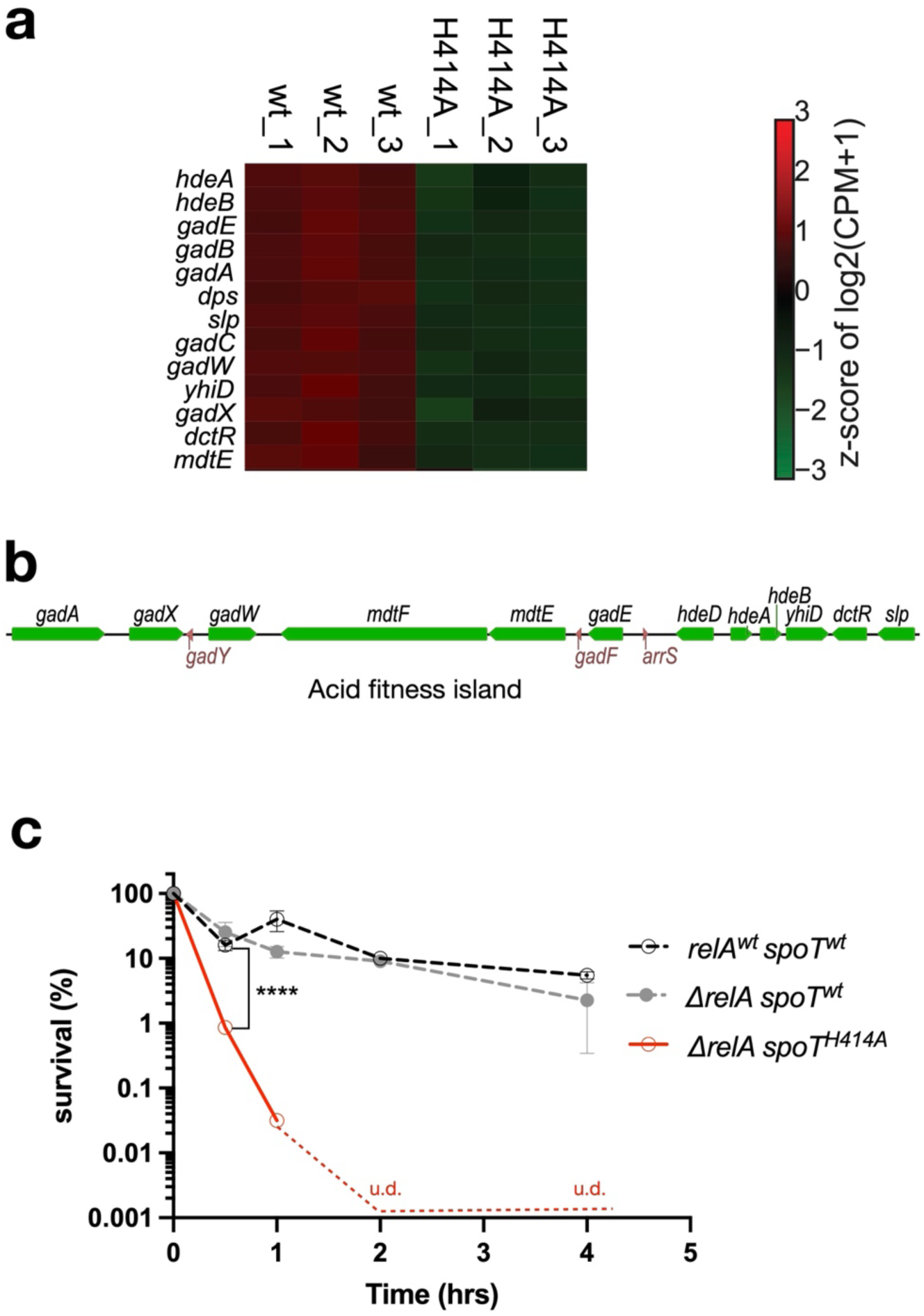
Transcriptomic analysis reveals collapse of acid resistance programs. **(a)** Heatmap of genes encoded in the *E. coli* acid fitness island in *spoT^H414A^* relative to *spoT^wt^*. **(b)** Genetic organization of the *E. coli* acid fitness island. (**c**) Acid resistance assay of wild type MG1655 (*relA*^wt^ *spoT^wt^*), *ΔrelA spoT^H414A^* and *ΔrelA spoT^wt^*. Survival of indicated strains following challenge at pH 2.5, expressed as the percentage of survival (i.e., colony-forming units (CFUs) remaining relative to starting CFUs) for each strain. Data are means ± s.d. from biological replicates. u.d., undetectable. Statistical significance was assessed using unpaired two-tailed Student’s *t*-tests. **** denote *P* < 0.0001.

### SpoT, but not RelA, is required for survival under extreme acid stress

Guided by the pronounced transcriptional repression of acid resistance genes in *spoT^H414A^* cells, we next examined survival under lethal acid stress. Upon exposure to pH2.5, *spoT^H414A^* cells exhibited profound acid sensitivity, with viability declining rapidly and no detectable survivors after 60 minutes (**Fig. 2c**). In contrast, both the parental wild-type strain and the *spoT^wt^* strain survived acid challenge efficiently, indicating that RelA is dispensable for acid resistance under this condition.

Notably, despite the severe acid sensitivity of *spoT^H414A^* cells, viability under neutral, well-buffered conditions (M9Glc) was preserved (**Extended Data Fig. 1g**), suggesting that SpoT H414 is specifically required to support stress preparedness rather than basal cell viability. Together, these results identify SpoT, rather than RelA, is required to support survival under extreme acid stress in *E. coli*.

### Loss of SpoT regulation drives arginine overflow and glutamate depletion

To define the metabolic basis of the growth defect observed in *spoT^H414A^*cells, we examined transcriptomic changes in central metabolic pathways. While expression of many amino acid biosynthetic genes was modestly reduced, one pathway was prominently induced: arginine biosynthesis was strongly and coordinately upregulated (**Fig. 3a, 3b**). All the nine enzymes of the arginine biosynthetic pathway exhibited significantly increased expression, together with *carAB*, which supplies carbamoylphosphate to this pathway. This pattern indicates a pronounced redirection of metabolic flux toward arginine production.

**Figure 3.**
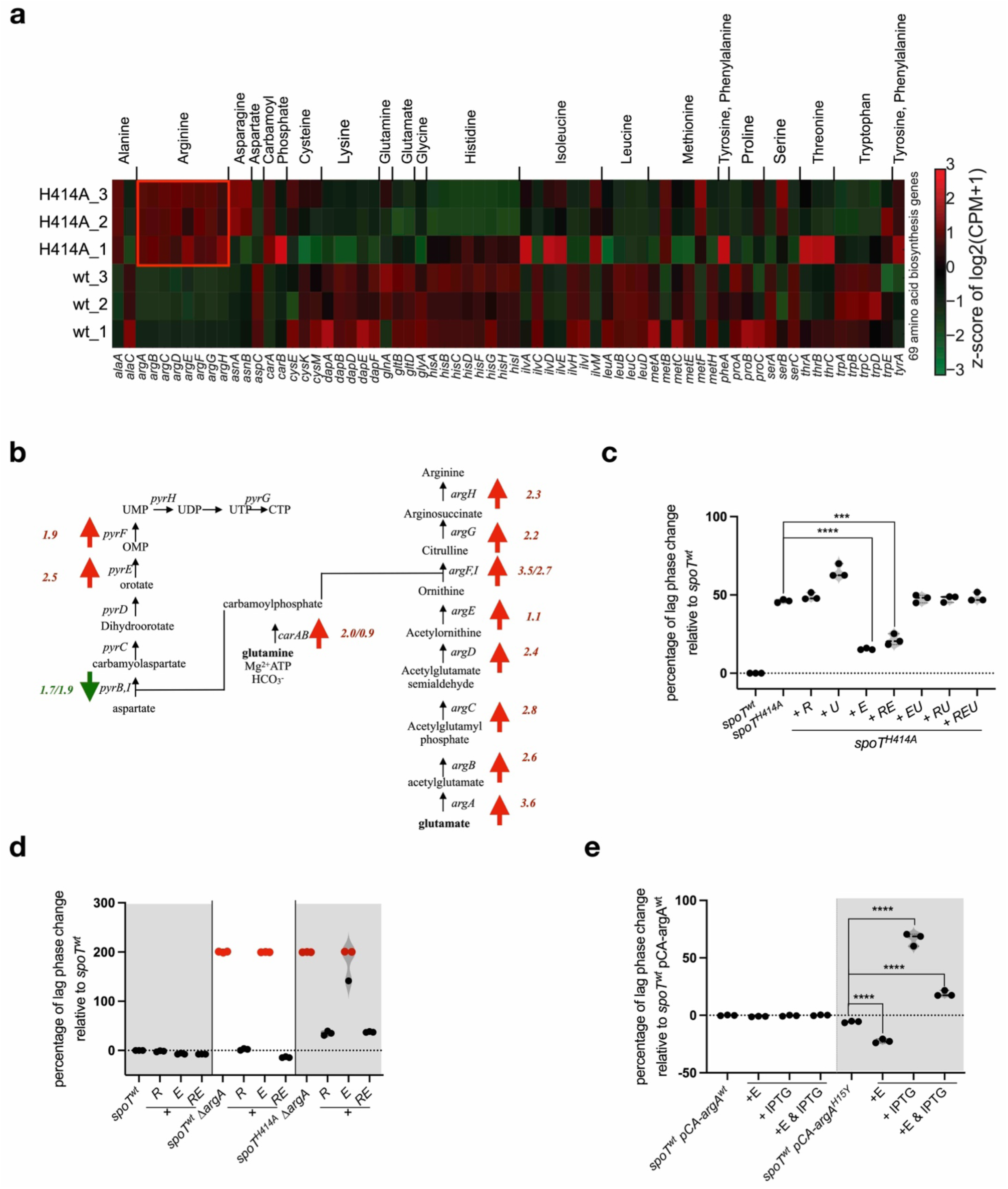
Arginine metabolic overflow drives glutamate depletion in *spoT^H414A^*. **(a)** Heatmap of genes involved in amino acid biosynthesis, comparing *spoT^H414A^* and *spoT^wt^*strains. The arginine biosynthesis genes are highlighted. **(b)** Schematic of the arginine biosynthesis pathway in *E. coli*. Genes upregulated in *spoT^H414A^*cells are indicated by red upward arrows, accompanied by average log₂ fold-change values relative to *spoT^wt^*. **(c)** Normalized lag phase duration of *spoT^H414A^*cells grown in M9Glc medium with supplementation of arginine (5.2 mM, R), uracil (0.2 mM, U), glutamate (0.6 mM, E), or the indicated combinations, expressed relative to the *spoT^wt^* strain. **(d)** Normalized lag phase duration of indicated strains grown in M9Glc medium with supplementation of arginine (R), glutamate (E), or both, relative to *spoT^wt^*. Red dots indicate failure to resume growth within the 72-h observation window. **(e)** Normalized lag phase duration of *spoT^wt^* strains harboring pCA24N-*argA^wt^* or the feedback-resistant pCA24N- *argA^H15Y^*allele, grown with or without IPTG induction and/or glutamate supplementation (0.6 mM), relative to the uninduced *argA^wt^* control. Data are means ± s.d. from biological replicates. Statistical significance was assessed using unpaired two-tailed Student’s *t*-tests. ****, *** denote *P* < 0.0001 and 0.001, respectively.

In contrast, genes directing carbamoylphosphate toward pyrimidine nucleotide biosynthesis were downregulated, suggesting selective diversion of this shared intermediate rather than a generalized increase in anabolic metabolism. Because arginine biosynthesis consumes glutamate as a precursor, the coordinated induction of the *arg* operons is predicted to substantially deplete intracellular glutamate pools.

In parallel with arginine pathway induction, genes involved in *de novo* purine nucleotide biosynthesis were broadly upregulated (**Extended Data Fig. 2**). Many of these reactions consume glutamine as a nitrogen donor, which is regenerated from glutamate, thereby imposing an additional, indirect demand on glutamate availability. Thus, the combined activation of arginine and purine biosynthetic programs is expected to exacerbate glutamate depletion, compounding the metabolic imbalance caused by SpoT H414A mutation.

Together, these transcriptional signatures point to a misallocation of metabolic resources upon loss of SpoT H414, with excessive flux into arginine biosynthesis and increased purine nucleotide production collectively depleting glutamate. Given the central role of glutamate in both metabolism and acid resistance, this imbalance offers a unifying explanation for the growth and stress survival defects observed in *spoT^H414A^* cells and motivated direct tests of glutamate availability in subsequent experiments.

### Glutamate depletion contributes to growth defects caused by SpoT perturbation

Prompted by the transcriptomic evidence for glutamate depletion, we tested whether exogenous supplementation could restore growth. Addition of glutamate to M9Glc robustly improved growth of *spoT^H414A^* cells, substantially shortening the lag phase and increasing final biomass (**Fig. 3c**). In contrast, supplementation with arginine or uracil failed to restore growth, and co-supplementation with glutamate conferred no additional benefit.

Among all 20 amino acids, supplementation with glutamine, asparagine, or proline partially restored growth, consistent with their metabolic proximity to glutamate, but none matched the effect of glutamate itself (**Extended Data Fig. 3a**). Increasing the concentration of glutamate up to tenfold failed to further improve growth (**Extended Data Fig. 3b**), and combining glutamate with glutamine, asparagine, or proline did not enhance regrowth beyond that achieved with glutamate alone (**Extended Data Fig. 3c**). In contrast, supplementation with all 20 amino acids fully restored the growth defect of *spoT^H414A^* cells to a level exceeding that of the *spoT^wt^* strain (**Extended Data Fig. 3d**), indicating that glutamate depletion represents a dominant, though not exclusive, metabolic limitation associated with SpoT H414A dysfunction. Importantly, glutamate supplementation had only minimal effects on the growth of a ppGpp⁰ (*ΔrelA ΔspoT*) strain (**Extended Data Fig. 3e**), indicating that growth rescue requires SpoT-dependent basal ppGpp rather than nonspecific nutritional supplementation.

To directly test whether arginine biosynthesis drives glutamate depletion, we deleted *argA*, encoding the first committed enzyme of the arginine biosynthetic pathway. As expected, the resulting *spoT^wt^ ΔargA* strain failed to grow in M9Glc; this defect was fully rescued by arginine supplementation but not by glutamate, confirming that growth limitation in this background reflects arginine auxotrophy rather than glutamate deficiency (**Fig. 3d**). Similarly, the *spoT^H414A^ ΔargA* strain failed to grow in M9Glc (**Fig. 3d**). Notably, arginine, but not glutamate, largely restored growth in this background, indicating that blocking arginine biosynthesis prevents excessive glutamate consumption and alleviates the growth defect of *spoT^H414A^*.

Arginine biosynthesis is normally subject to feedback inhibition through ArgA. Conversely, we found that artificial overexpression of a feedback-resistant ArgA variant (ArgA^H15Y^)^37^ via IPTG induction phenocopied the *spoT^H414A^* growth defect in M9Glc, and this delayed growth was substantially alleviated by glutamate supplementation (**Fig. 3e**). Together, these results establish that SpoT H414 restrains arginine-driven glutamate consumption, thereby preserving amino acid homeostasis required for growth in minimal medium.

### SpoT perturbation compromises glutamate-dependent acid resistance

Because glutamate is the essential substrate for the Gad acid resistance system, we next asked whether glutamate depletion contributes to the acid sensitivity of *spoT^H414A^* cells. Supplementation of the acid challenge medium with glutamate fully rescued survival during short-term acid exposure and markedly improved long-term viability (**Fig. 4a**). In contrast, supplementation with arginine or lysine provided only modest protection (**Extended Data Fig. 4a**), consistent with the prominent contribution of the glutamate-dependent Gad system under extreme acid stress.

**Figure 4.**
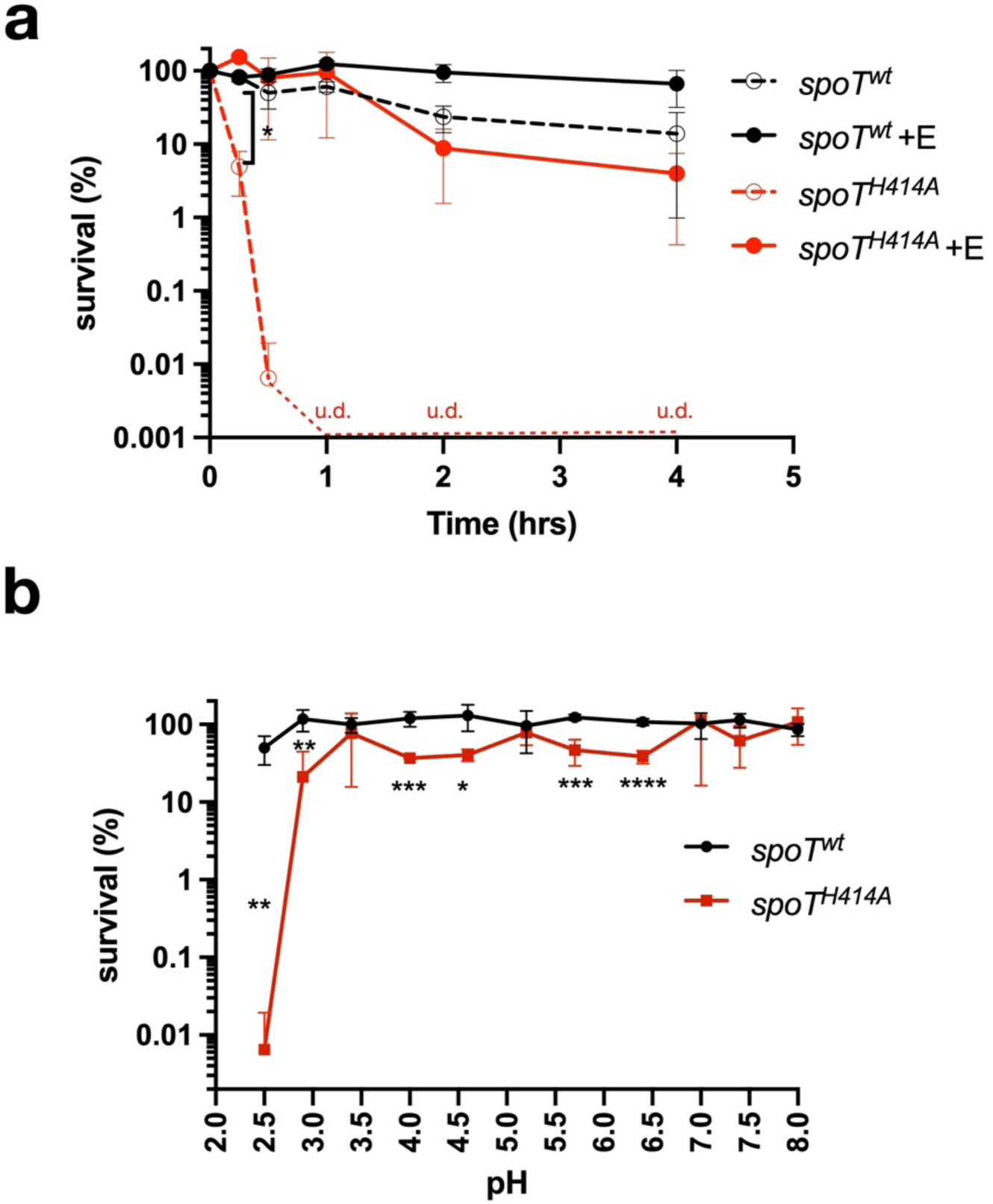
SpoT H414 is required for survival under extreme acid stress. **(a)** Acid resistance assay of *ΔrelA spoT^H414A^* and *ΔrelA spoT^wt^* as in Figure 2c. Where indicated (+E), glutamate (0.6 mM) was included in the acid challenge medium. u.d., undetectable. **(b)** Survival of *spoT^H414A^* and *spoT^wt^*strains following 30 min challenge in buffers adjusted to the indicated pH values, normalized to starting CFUs. Data are means ± s.d. from biological replicates. Statistical significance was assessed using unpaired two-tailed Student’s *t*-tests. ****, ***, **, * denote *P* < 0.0001, 0.001, 0.01, and 0.05, respectively.

To determine whether impaired regulation of the Gad system accounted for the acid resistance defect, we ectopically expressed *gadE*, the master transcriptional activator of Gad genes. Even low-level expression of *gadE* significantly restored acid resistance in *spoT^H414A^*cells, and combined supplementation with glutamate further enhanced survival (**Extended Data Fig 4b**). These interventions did not alter acid resistance in *spoT^wt^* cells, indicating that the defect arises specifically from SpoT dysfunction. Notably, despite its ability to restore acid survival, ectopic expression of *gadE* failed to improve the growth of *spoT^H414A^* cells in M9Glc (**Extended Data Fig. 4c**). This functional separation indicates that impaired acid resistance is not sufficient to account for the growth defect, which is more closely associated with glutamate depletion and broader metabolic misallocation.

We further examined acid tolerance across a range of pH values. In addition to catastrophic sensitivity at pH 2.5, *spoT^H414A^* cells exhibited reduced survival under mildly acidic conditions (pH 4.0-6.0) (**Fig. 4b, Extended Data Fig 4d**), consistent with a broader impairment of multiple acid stress response pathways (**Extended Data Fig 4e**). Notably, these defects occurred despite preserved viability under neutral pH conditions (**Extended Data Fig 1g**), indicating that SpoT-mediated basal regulation selectively primes cells for acid stress rather than broadly supporting growth.

### Intragenic suppression reveals intrinsic regulation within SpoT

These findings raised a fundamental question of how SpoT activity is regulated to maintain the physiological state underlying growth and stress resistance. The severe growth and acid resistance defects of *spoT^H414A^*cells suggested that H414 plays a central regulatory role in SpoT function. To determine whether this phenotype reflected impaired regulation by trans-acting protein factors, we first tested overexpression of established SpoT regulators, including ObgE^38^, Rsd^39^, YtfK^40^, and AcpP^25^, none of which measurably restored growth of *spoT^H414A^* cells in minimal medium (**Extended Data Fig 5**). We further explored the possibility that H414 mediates regulation through interaction with previously unidentified protein partners by introducing a photo-crosslinkable unnatural amino acid at position 414 and performing UV-activated crosslinking under the same growth conditions. Despite robust expression of full-length SpoT proteins, we did not detect reproducible interactions with additional protein factors (**Extended Data Fig. 6**). Together, these results suggested that the regulatory function of H414 is unlikely to be mediated by trans-acting protein partners. We therefore pursued an unbiased genetic approach and isolated spontaneous suppressor mutants capable of restoring growth of *spoT^H414A^* cells on M9Glc. After prolonged incubation (24-72 h), colonies of varying sizes emerged on M9Glc agar (**Fig. 5a**).

**Figure 5.**
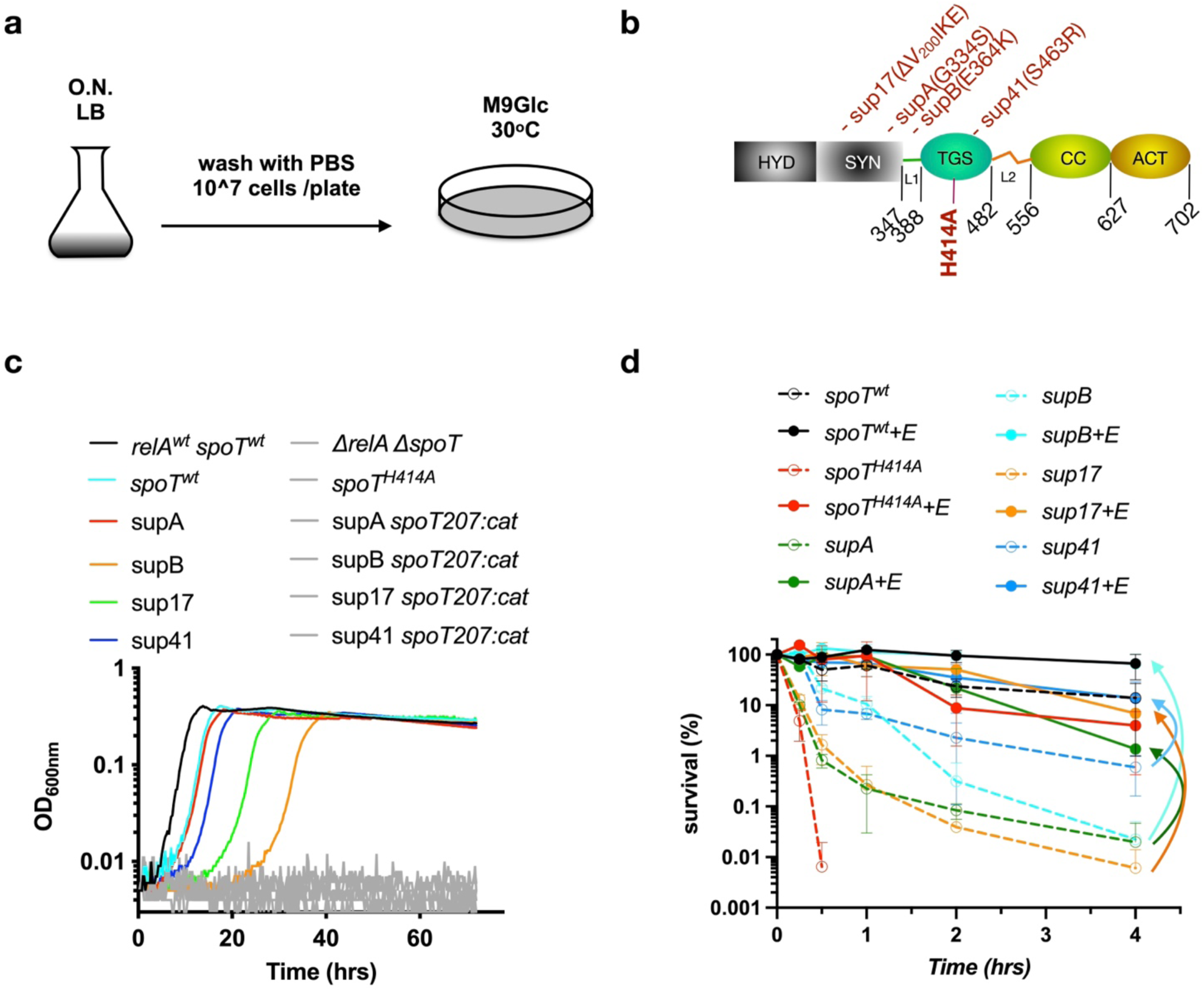
Intragenic SpoT suppressor mutations restore growth and acid resistance. **(a)** Schematic illustrating the isolation of spontaneous suppressor mutations that enable growth of *spoT^H414A^* cells on M9Glc medium. **(b)** Domain architecture of the SpoT protein, indicating the locations of suppressor mutations. Amino-acid positions are shown below. L1/L2, linker-1/linker-2; HYD, hydrolase domain; SYN, synthetase domain; TGS, ThrRS-GTPase-SpoT domain; CC, conserved cysteine domain; ACT, aspartate kinase-chorismate mutase-TyrA domain. **(c)** Growth dynamics of four suppressor strains (supA, supB, sup17, sup41) compared with *spoT^H414A^*, *spoT^wt^*, ppGpp⁰ (*ΔrelA ΔspoT*), and wild-type MG1655 (*relA^wt^ spoT^wt^*). Also shown are suppressor strains in which *spoT* was deleted (*spoT207::cat*). **(d)** Survival of indicated strains following challenge at pH 2.5. Where indicated (+E), glutamate (0.6 mM) was included. Data are means ± s.d. Statistical significance was assessed using unpaired two-tailed Student’s *t*-tests. ***, **, * denote *P* < 0.001, 0.01, and 0.05, respectively.

Analysis of 50 independently isolated suppressors revealed that the strongest suppressors predominantly carried secondary mutations within the *spoT* gene itself. Among large-colony suppressors, we identified recurrent substitutions at residues G334 (G334S; supA, 17/50), E364 (E364K; supB, 5/50), and S463 (S463R; sup41, 9/50) as well as a small in-frame deletion within the synthetase domain (ΔV_200_IKE; sup17, 3/50) (**Fig. 5b**). Whole-genome sequencing of small-colony suppressors lacking *spoT* mutations identified a single mutation in *rpoB* (K1262Q), which is likely similar as the previously described stringent-response suppressors^41^ supporting the growth of ppGpp^0^ strain on minimal media, and therefore these strains were excluded from further analysis.

Reconstruction of the identified *spoT* suppressor mutations in a clean background fully recapitulated growth restoration of *spoT^H414A^* in M9Glc (**Extended Data Fig 7**), whereas deletion of *spoT* in the suppressor strains abolished growth (**Fig. 5c**). These results demonstrate that intragenic mutations within SpoT are both necessary and sufficient to suppress the *spoT^H414A^* growth defect.

We next asked whether suppression of the growth defect was accompanied by restoration of acid resistance. All four *spoT* suppressor strains exhibited improved survival under lethal acid challenge (pH 2.5) compared to the parental *spoT^H414A^* strain, although the degree of rescue varied (**Fig. 5d**). The suppressor sup41 displayed the most robust acid resistance, followed by supB, supA and sup17. Supplementation of the acid challenge medium with glutamate further enhanced survival across all suppressor strains, with particularly pronounced effects in the weaker suppressors supB and sup17 (**Fig. 5d**). These differences suggest that individual suppressor mutations restore SpoT function through distinct mechanisms that differentially affect glutamate homeostasis and stress preparedness.

Together, these results show that intragenic suppression of SpoT H414A simultaneously restores growth and acid resistance, reinforcing the link between SpoT activity, metabolic balance, and stress survival.

### Basal ppGpp is actively maintained within a narrow functional window

Given SpoT’s dual role as a ppGpp synthase and hydrolase, we next asked whether the growth defect of *spoT^H414A^* cells is associated with altered ppGpp homeostasis, and whether suppressor mutations restore a physiologically relevant ppGpp state. To address this, we measured intracellular ppGpp levels using ^32^P-orthophosphate labeling and thin-layer chromatography^18^. As expected, *spoT*^wt^ cells produced a readily detectable basal ppGpp signal, whereas *spoT^H414A^* cells accumulated only marginal, near-background levels of ppGpp (**Fig. 6a**). Notably, ppGpp levels in the suppressor strains remained close to the detection limit and were not substantially elevated relative to *spoT^H414A^*cells.

**Figure 6.**
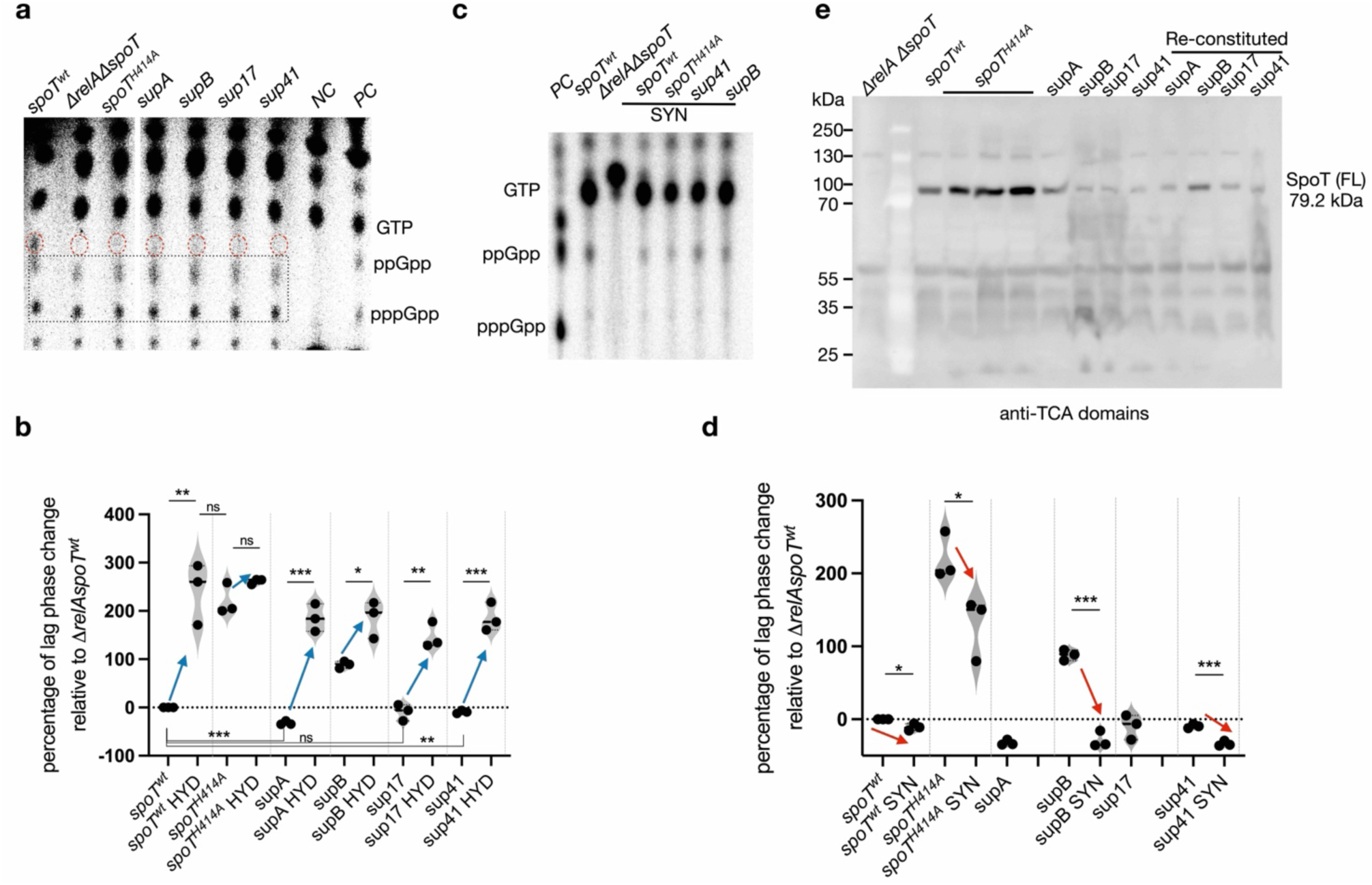
A sub-basal ppGpp pool is sufficient for physiological function. **(a)** Thin-layer chromatography (TLC) analysis of ^32^p-labeled nucleotides from indicated strains grown in low-phosphate MOPS minimal medium supplemented with 0.2% glucose. NC and PC denote valine-treated *ΔrelA ΔspoT* and wild-type MG1655 controls, respectively. Red circles mark ppGpp; boxed regions indicate nonspecific spots also present in *ΔrelA ΔspoT* samples. **(b)** Effect of inactivating SpoT ppGpp synthetase activity (E319Q; HYD) on growth recovery in M9Glc. Lag phase duration is normalized to *spoT^wt^*. **(c)** TLC analysis of ^32^p-labeled nucleotides from strains carrying SpoT hydrolase-inactive mutations (H72A D73A; SYN). **(d)** Growth recovery of hydrolase-inactive strains (SYN), quantified as normalized lag phase duration relative to *spoT^wt^*. Note that the hydrolase-inactive (SYN) variants of supA and sup17 could not be obtained, thereby not tested. **(e)** Immunoblot analysis of SpoT protein levels using an antiserum raised against the SpoT C-terminal (TGS-CC-ACT; anti-TCA) region. Cells were grown in M9Glc to early exponential phase (OD_600nm_ ≈ 0.2) prior to sampling. *spoT^H414A^* samples were collected after prolonged incubation, as in Extended Data Fig. 1c. *ΔrelA ΔspoT* cells grown in LB served as a negative control.

This observation raised the possibility that either very small increases in ppGpp are sufficient to restore physiological function, or that ppGpp turnover is highly dynamic in suppressor strains. To distinguish between these possibilities, we genetically perturbed SpoT enzymatic activities. Inactivation of the ppGpp synthetase activity of SpoT by the E319Q substitution in suppressor backgrounds severely delayed or abolished growth recovery in M9Glc (**Fig. 6b**), demonstrating that ppGpp synthesis is required for suppression. Conversely, inactivation of SpoT’s hydrolase activity by the H72A/D73A substitution led to robust ppGpp accumulation (**Fig. 6c**) and accelerated growth resumption (**Fig. 6d**), indicating that suppressor strains produce ppGpp that is rapidly turned over under basal conditions. Notably, the hydrolase-inactive variants of supA and sup17 could not be obtained, likely due to the toxicity associated with excessive ppGpp accumulation.

Because C-terminal domains of RelA/SpoT homologs are known to regulate the catalytic activities of the N-terminal enzymatic domains, we next systematically truncated these regulatory regions to assess how suppressor mutations alter intramolecular control of SpoT activity. Consistent with this model, truncation of individual C-terminal domains revealed that distinct suppressor mutations differentially modify the contribution of these domains to growth control (**Extended Data Fig. 8**), without abolishing the requirement for basal ppGpp synthesis. These findings indicate that SpoT H414 functions within a broader intramolecular regulatory network that fine-tunes basal ppGpp production, rather than acting as an isolated molecular switch.

Together, these data demonstrate that a sub-basal but non-zero ppGpp pool - close to the detection limit of conventional assays - is sufficient to support growth in minimal medium, and that SpoT activity is finely tuned to maintain this pool within a narrow physiological range.

### Negative feedback between ppGpp and SpoT abundance stabilizes basal ppGpp

To investigate how SpoT activity is regulated, we examined SpoT protein levels using an antiserum raised against the C-terminal regulatory domains of SpoT. Immunoblot analysis revealed that SpoT protein levels were markedly elevated in *spoT^H414A^* cells compared to *spoT*^wt^ and suppressor strains (**Fig. 6e**).

Because *spoT^H414A^* cells produce sub-basal ppGpp levels, whereas suppressor strains restore physiological function with slightly elevated but tightly controlled ppGpp, these results suggest the existence of a negative feedback mechanism in which basal ppGpp limits SpoT abundance. Such feedback would stabilize intracellular ppGpp levels by adjusting SpoT protein concentration in response to alarmone availability. This regulatory relationship provides a parsimonious mechanistic framework for how SpoT may maintain a narrow basal ppGpp range sufficient for metabolic homeostasis without eliciting an acute stringent response.

### Graded restoration of transcriptional programs reflects quantitative tuning of basal SpoT activity

To understand how suppressor mutations restore growth and acid resistance, we performed RNA sequencing on the four suppressor strains. Parallel coordinates and multidimensional scaling analyses revealed that strong suppressors (supA and sup41) clustered closely with *spoT*^wt^ cells, whereas weaker suppressors (supB and sup17) occupied intermediate positions between *spoT^H414A^* and *spoT*^wt^ (**Extended Data Fig. 9a,9b**).

Across all suppressors, a core set of genes involved in amino acid metabolism and stress adaptation was consistently restored (**Extended Data Fig. 9c**), whereas genes aberrantly induced in *spoT^H414A^* cells were broadly repressed (**Extended Data Fig. 9d**). Moreover, strong suppressors (supA and sup41) largely restored expression of acid resistance genes and concomitantly repressed phage shock and SOS response pathways; in contrast, weaker suppressors (supB and sup17) exhibited only partial correction of these transcriptional defects (**Extended Data Fig. 10**), consistent with their physiological phenotypes (**Fig. 5c**).

Together, these results indicate that suppressor mutations restore SpoT-dependent regulation in a graded fashion, with transcriptional recovery closely correlating with physiological rescue.

### The regulatory function of SpoT H414 is conserved across enteric pathogens

Finally, we asked whether the regulatory role of SpoT H414 is conserved in pathogenic enterobacteria. We introduced the corresponding H414A substitution into SpoT homologs from *Salmonella enterica* serovar Typhimurium and *Shigella flexneri* and expressed these proteins in an *E. coli* ppGpp⁰ background.

Wild-type SpoT homologs from both species largely restored growth in M9Glc and acid resistance, whereas the corresponding H414A variants failed to support either phenotype (**Fig. 7a-d**). Notably, glutamate supplementation rescued both growth and acid resistance defects in the *Shigella* SpoT_sf_ H414A strain but not in the *Salmonella* SpoT_st_ H414A variant. This differential rescue correlated with the extent of sequence divergence within the C-terminal regulatory regions of SpoT. *S. flexneri* SpoT_sf_ differs from *E. coli* SpoT_ec_ at only a single conserved position (A614G) within the CC domain, whereas *S. Typhimurium* SpoT_st_ shares 98.15% overall identity but contains multiple amino acid substitutions distributed across the C-terminal domains, including a cluster of nine substitutions within linker-2 between TGS and CC domains (**Extended Data Fig. 11**). These sequence differences are likely to modulate intramolecular regulatory interactions within SpoT, thereby influencing the ability of the H414A variants to respond to glutamate supplementation, akin to the differential behaviors of the intragenetic suppressor strains (**Fig. 5d**).

**Figure 7.**
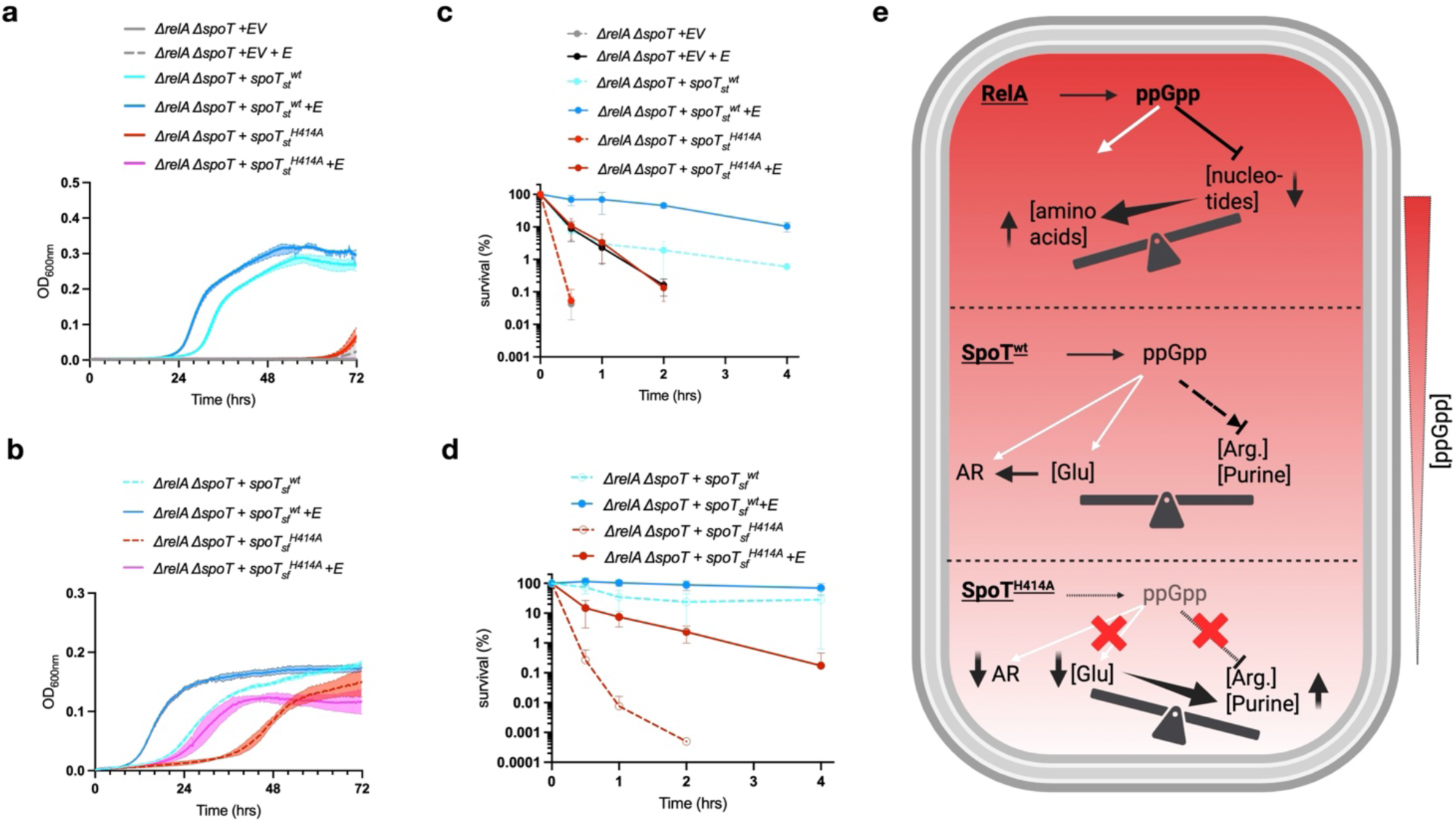
Conservation of SpoT H414 function and model for basal ppGpp regulation. **(a,b)** Growth of *E. coli* ppGpp⁰ strains complemented with SpoT homologs from *Salmonella enterica* serovar Typhimurium (SpoT_st_) (**a**) or *Shigella flexneri* (SpoT_sf_) (**b**). E, glutamate (0.6 mM); EV, empty vector. Data represent at least three biological replicates. **(c,d)** Survival of complemented ppGpp⁰ strains expressing SpoT_st_ (**c**) or SpoT_sf_ (**d**)) following challenge at pH 2.5, normalized to starting CFUs. Where indicated (+E), glutamate (0.6 mM) was included. **(e)** Top: Under acute amino-acid starvation, RelA is rapidly activated by ribosome-associated signals to generate a high-amplitude ppGpp surge, globally reprogramming transcription and metabolism to suppress nucleotide pathways and promote amino-acid availability. Middle: SpoT maintains a low but physiologically essential basal ppGpp pool that is distinct from and independent of RelA. This basal ppGpp state coordinates metabolic homeostasis by restraining excessive arginine and purine biosynthesis, preserving intracellular glutamate levels, and priming multiple acid resistance programs, including the glutamate-dependent Gad system (AR). Bottom: In *spoT^H414A^* cells, a sub-basal level of ppGpp remains below the threshold required for effective metabolic and stress control. Therefore, arginine and purine biosynthetic pathways become overactivated, intracellular glutamate is depleted, and acid resistance programs are inadequately induced, resulting in growth failure in minimal medium and extreme sensitivity to acid stress.

Altogether, these results demonstrate that SpoT H414 plays a conserved and functionally important role in coordinating growth and acid resistance across enteric pathogens, highlighting the physiological and evolutionary relevance of basal SpoT regulation.

## Discussion

### Basal ppGpp defines a distinct regulatory regime coordinating metabolism and stress resistance

The stringent response has classically been viewed as an acute adaptation to amino acid starvation, driven by RelA-dependent bursts of ppGpp that globally reprogram transcription and metabolism (**Fig. 7e**, top). In this canonical framework, ppGpp functions as an emergency alarmone that rapidly suppresses growth-associated biosynthetic programs and redirects cellular resources upon severe nutrient limitation.

Our findings reveal a complementary and previously underappreciated mode of ppGpp signalling that operates outside this acute starvation paradigm. By dissecting the function of a conserved regulatory residue in SpoT, H414, we identify a basal ppGpp state that is actively maintained during nutrient-limited growth and is essential for coordinating metabolic homeostasis with stress preparedness (**Fig. 7e**, middle). Perturbation of this state uncovers a sharp physiological threshold below which metabolic balance and stress resistance collapse (**Fig. 7e**, bottom).

Importantly, the phenotypes observed in *spoT^H414A^* cells are not readily explained by defects in canonical RelA-mediated stringent response activation. All experiments were conducted in a *ΔrelA* background and under conditions that do not elicit acute amino acid starvation. Moreover, the transcriptional profile of *spoT^H414A^* cells is fundamentally distinct from those reported in classical stringent response studies driven by RelA-dependent ppGpp bursts^15,16^. Loss of SpoT H414 does not recapitulate either RelA-activated or ppGpp⁰ transcriptional states. Instead, *spoT^H414A^* cells exhibit a selective and imbalanced response marked by overcommitment to arginine and nucleotide biosynthesis, depletion of central metabolites such as glutamate, and failure to sustain specific stress-response programs, most notably the glutamate-dependent acid resistance system.

Together, these observations support the existence of a regulatory regime operating at ppGpp levels above zero yet below the basal state normally maintained by wild-type SpoT. Within this low-amplitude range, ppGpp does not function as a global transcriptional on-off switch, but instead acts as a quantitative tuning signal that constrains metabolic flux allocation and preserves stress preparedness. Loss of this basal control therefore exposes physiological vulnerabilities that are not captured by classical models of the stringent response. In this framework, SpoT functions not merely as a modulator of RelA activity, but as a primary regulator of a stabilized physiological state. This low but non-zero ppGpp levels act not as an emergency brake on growth, but as a buffering signal that integrates metabolism with stress preparedness during nutrient-limited growth.

### SpoT-mediated control of glutamate homeostasis underlies stress preparedness without compromising basal viability

A central mechanistic insight from this study is that SpoT-dependent basal ppGpp constrains metabolic flux allocation to preserve glutamate homeostasis. Transcriptomic profiling of *spoT^H414A^*cells revealed coordinated induction of arginine biosynthesis (**Fig. 3a,b**), which directly consumes glutamate, together with upregulation of *de novo* purine nucleotide biosynthesis (**Extended Data Fig. 3a,b**), which indirectly increases glutamate demand through enhanced glutamine utilization. This pattern reflects a loss of quantitative control over anabolic commitment rather than selective activation of a single biosynthetic pathway.

Genetic and physiological analyses identify arginine biosynthesis as the primary driver of glutamate depletion in this context. Blocking the first committed step of arginine synthesis prevents glutamate starvation and restores growth (**Fig. 3d**), whereas forced arginine overproduction phenocopies the *spoT^H414A^* defect (**Fig. 3e**).

Glutamate occupies a uniquely central position in bacterial physiology, serving both as a metabolic hub and as the essential substrate for the glutamate-dependent Gad acid resistance system. Accordingly, glutamate depletion provides a parsimonious explanation for the coupled phenotypes observed in *spoT^H414A^* cells, including impaired growth in minimal medium and failure of acid resistance. Glutamate supplementation and GadE-dependent activation of the acid resistance program rescue survival under low-pH challenge (**Fig. 4a; Extended Data Fig. 4b**), indicating that SpoT-dependent basal ppGpp enforces a metabolic prioritization strategy that preserves metabolites required for effective stress defence.

Importantly, disruption of this basal regulatory state does not cause immediate loss of viability under neutral, well-buffered conditions (**Extended Data Fig. 1g)**, despite severe repression of acid resistance genes and extreme sensitivity to low pH (**Fig. 2c**). This separation between basal growth capacity and stress survival indicates that SpoT-derived basal ppGpp primarily maintains physiological preparedness rather than minimal viability. This distinction helps explain why the most pronounced phenotypes of *spoT^H414A^* cells emerge only upon environmental challenge.

Notably, RelA is dispensable for acid resistance under the conditions examined (**Fig. 2c**), highlighting a functional division of labour between the two stringent response enzymes. Whereas RelA mediates acute, high-amplitude responses to amino acid starvation, SpoT maintains a low-level regulatory state that buffers metabolic allocation and supports long-term stress readiness.

### SpoT maintains basal ppGpp through coupled activity tuning and abundance feedback

The identification of intragenic suppressors of *spoT^H414A^*reveals that SpoT encodes an intrinsic regulatory system capable of quantitatively tuning basal ppGpp levels. Suppressor mutations distributed across both the catalytic and regulatory regions of SpoT restore growth (**Fig. 5c**) and acid resistance (**Fig. 5d**) without producing large, readily detectable increases in ppGpp (**Fig. 6a**). These observations indicate that physiological rescue does not require restoration of wild-type ppGpp abundance, but instead depends on re-establishing a narrowly defined sub-basal ppGpp state, consistent with the graded restoration of transcriptional programs observed across suppressor strains (**Extended Data Fig. 9,10**).

Direct perturbation of SpoT enzymatic activities supports this interpretation. Inactivation of ppGpp synthesis abolishes suppressor-mediated growth recovery (**Fig. 6b**), whereas inactivation of SpoT hydrolase activity leads to substantial ppGpp accumulation (**Fig. 6c**) and accelerated growth resumption (**Fig. 6d**). These results indicate that suppressor mutations rebalance SpoT synthetic and hydrolytic activities rather than bypassing the requirement for ppGpp. The inability to construct synthetase-only variants in specific suppressor backgrounds further underscores the importance of quantitative control. In these contexts, elevated ppGpp levels are likely deleterious, consistent with the well-established growth-inhibitory effects of excessive ppGpp. Thus, both insufficient and excessive ppGpp are incompatible with optimal growth, defining a sharply constrained physiological window for basal ppGpp function.

However, tuning of catalytic activity alone is unlikely to be sufficient to stabilize a low-amplitude signalling state over extended periods of growth. Consistent with this, SpoT protein levels are markedly elevated in *spoT^H414A^* cells but reduced in suppressor backgrounds (**Fig. 6e**), revealing an additional layer of regulation mediated by negative feedback on SpoT abundance. This inverse relationship suggests that basal ppGpp limits SpoT accumulation, thereby constraining total enzymatic capacity for ppGpp turnover.

Together, quantitative tuning of SpoT catalytic activity and feedback control of SpoT abundance may enable precise maintenance of ppGpp within a narrow functional window. This coupled regulatory architecture provides a mechanistic model for how bacteria stabilize a low-level ppGpp signalling state sufficient for metabolic coordination and stress preparedness, yet below the threshold that triggers growth arrest and the canonical stringent response.

### Conservation and implications of basal SpoT regulation

The functional conservation of H414 across *Escherichia coli*, *Salmonella*, and *Shigella* SpoT proteins demonstrates that this residue is broadly required for growth in minimal medium and survival under extreme acid stress (**Fig. 7a-d**). At the same time, the differential ability of *Salmonella* and *Shigella* SpoT H414A variants to respond to glutamate supplementation, akin to the intragenetic suppressors (**Fig. 5d**), suggests that sequence divergence within C-terminal regulatory regions modulates how basal SpoT regulation is implemented across species.

These observations indicate that, while the requirement for SpoT-mediated basal ppGpp control is conserved, individual bacteria may tune SpoT regulatory domains to optimize metabolic and stress-response coordination according to their metabolic capacity, ecological niches and life cycles. In this framework, basal ppGpp does not represent a rigid or uniform signalling state, but rather a conserved regulatory principle flexibly adapted through species-specific intramolecular tuning of SpoT.

Several important questions remain, including the nature of the signal sensed by SpoT H414 and the absolute intracellular concentrations and temporal dynamics of basal ppGpp. Addressing these questions will require new approaches capable of resolving low-abundance alarmone pools with high sensitivity, ideally at single-cell resolution, and across diverse growth conditions and bacterial species. Elucidating how basal ppGpp selectively modulates distinct metabolic and stress-response pathways will further clarify how this regulatory regime integrates diverse physiological demands.

Given the central role of acid resistance in enteric pathogenesis, SpoT-mediated basal ppGpp regulation may represent a previously underappreciated vulnerability that could be exploited to compromise bacterial survival under host-associated stresses.

## EXTENDED DATA FIGURE LEGENDS

**Extended Data Fig. 1.**
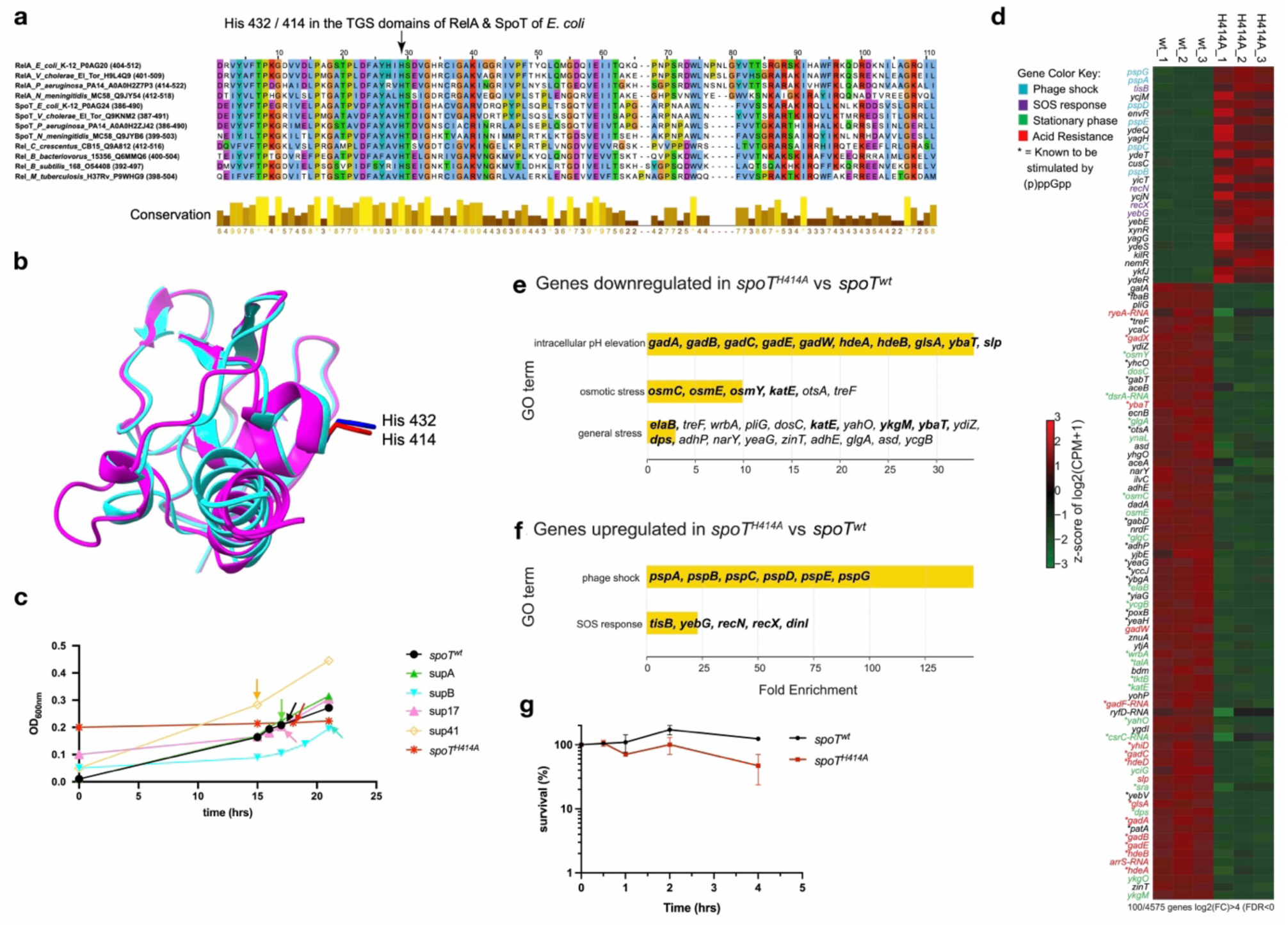
**(a)** Sequence alignment of 12 TGS domains of RelA, SpoT and their homologous proteins from α, β, γ and δ proteobacteria, Firmicute and Actinomycete. The highly conserved position corresponding to His 432 of RelA and 414 of SpoT of *E. coli* is indicated with an arrow. **(b)** Superpositioning of the TGS domains of RelA and SpoT from *E. coli* (rmsd over 91 of 98 atom pairs is 0.762 Å) generated by AF3. **(c)** Representative growth curves of *ΔrelA spoT^H414A^*, *ΔrelA spoT^wt^* and the four suppressor strains in M9Glc media for collecting RNA sequencing samples. Cells were collected when their OD_600nm_ approach ca. 0.2, except for *spoT^H414A^*cells, which could not grow in this medium, thus these cells were collected after incubated for 18 hrs in the same medium. Coloured arrows indicate the sampling points for each strain. **(d)** Heatmap of genes with greater than 16-fold changes between *ΔrelA spoT^H414A^* and *ΔrelA spoT^wt^* cells growing in M9Glc medium. **(e)** Gene ontology groups of genes downregulated in the *spoT^H414A^* strain. Genes downregulated at least 16-fold were extracted using DEGUST (|Log2(FC)| > 4; FDR < 0.05) and analysed by AmiGO and Panther. The 58 genes downregulated more than 16-fold (Log2(FC) < -4) are listed in **Table S3**. **(f)** Gene ontology groups of genes upregulated at least 16-fold in the *spoT^H414A^*strain. The 28 genes upregulated more than 16-fold (Log2(FC) > 4) are listed in **Table S3**. Note the differences in the Fold Enrichment scales in **(e)** and **(f)**. Gene names in bold in (**e, f**) are strongly up or downregulated. **(g)** Normalized percentage of surviving viable cells of the *spoT^wt^* and *spoT^H414A^*strains growing in M9Glc medium. Two replicates were performed and presented.

**Extended Data Fig. 2.**
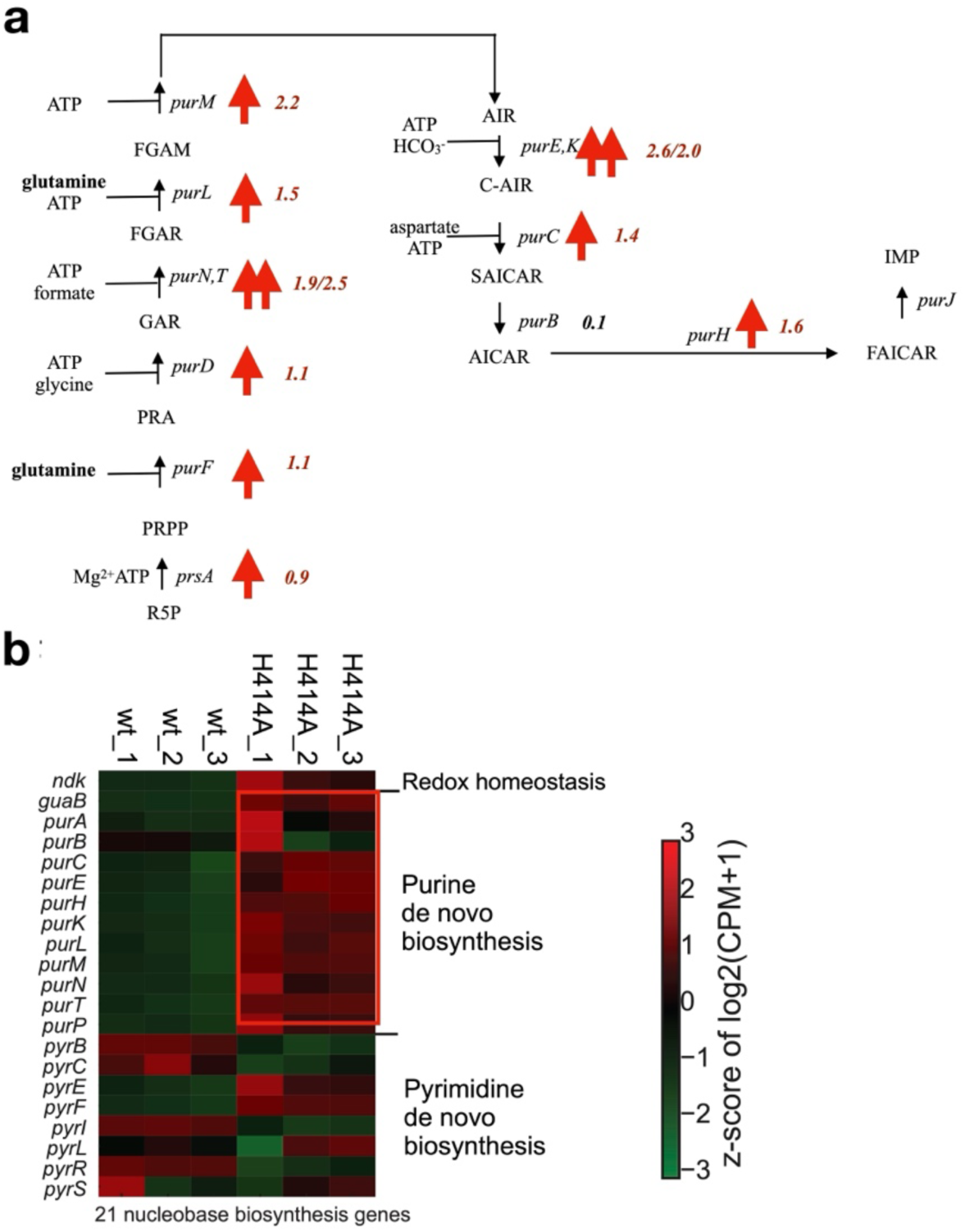
**(a)** Purine nucleotide de novo biosynthesis pathway in *E. coli*. The upregulated genes are indicated by a red upward arrow alongside average log2(FC) values of *spoT^H414A^*versus *spoT^wt^*. R5P, ribose-5-phosphate; PRPP, 5-phosphoribosyl 1-pyrophosphate; PRA, 5-phosphoribosylamine; GAR, 5’-phosphoribosylglycinamide; FGAR, 5’-phosphoribosyl-N-formylglycineamide; FGAM, 5’-phosphoribosylformylglycinamidine; AIR, 5-aminoimidazole ribonucleotide; C-AIR, 5-carboxyamino-1-(5-phospho-D-ribosyl)imidazole; SAICAR, 1-(5-phosphoribosyl)-4-(N-succino-carboxamide)-5-aminoimidazole; AICAR, 5-amino-1-(5-phospho-D-ribosyl)imidazole-4-carboxamide; FAICAR, 5-formamido-1-(5-phospho-D-ribosyl)imidazole-4-carboxamide; IMP, inosine 5’- monophosphate. **(b)** Heatmap of genes involved in purine and pyrimidine nucleotide de novo biosynthesis. The red box indicates purine de novo biosynthesis genes.

**Extended Data Fig. 3.**
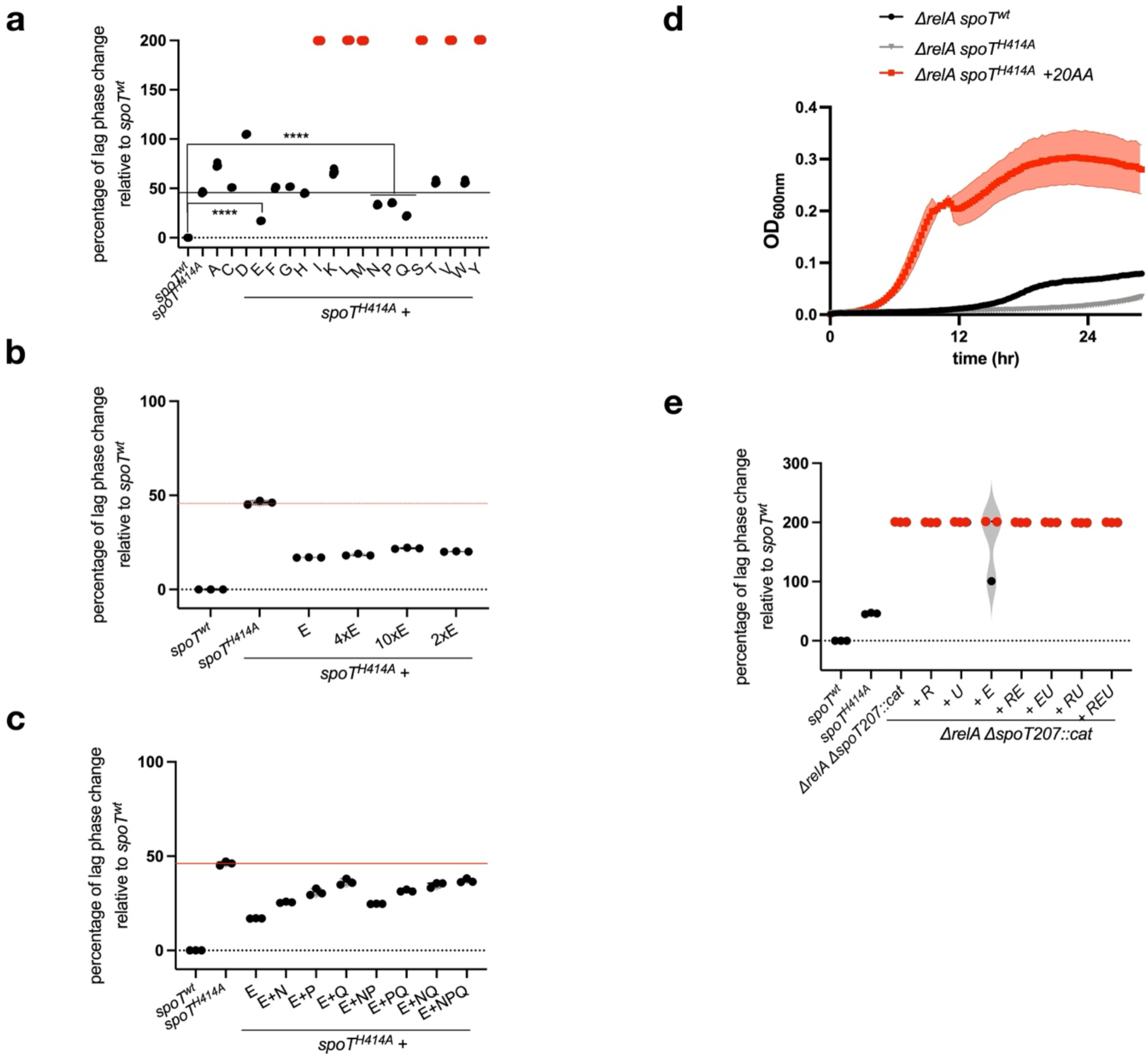
**(a)** Normalized lag phase length of the *spoT^H414A^* strain without or with supplemented each of the 20 amino acids (in single letters; A, alanine 0.8mM; C, cysteine 0.1mM; D, aspartic acid 0.4mM; E, glutamate 0.6mM; F, phenylalinine 0.4mM; G, glycine 0.8mM; H, histidine 0.2mM; I, isoleucine 0.4mM; K, lysine 0.4mM; L, leucine 0.8mM; M, methionine 0.2mM; N, asparigine 0.4mM; P, proline 0.4mM; Q, glutamine 0.6mM; R, arginine 5.2mM; S, serine 10mM; T, threonine 0.4mM; V, valine 0.6mM; W, tryptophan 0.1mM; Y, tyrosine 0.2mM), relative to the *spoT^wt^* strain in M9Glc medium. **(b)** Normalized lag phase length of the *spoT^H414A^* strain without or with supplemented 1x, 2x, 4x, 10x glutamate (E, 1x = 0.6 mM), relative to the *spoT^wt^* strain in M9Glc medium. **(c)** Normalized lag phase length of the *spoT^H414A^* strain without or with supplemented glutamate (0.6 mM) together with glutamine (Q, 0.6 mM), proline (P, 0.4 mM), asparagine (N, 0.4 mM), relative to the *spoT^wt^*strain in M9Glc medium. **(d)** Representative growth curves of *ΔrelA spoT^H414A^*, *ΔrelA spoT^wt^* strains without or with all 20 amino acids (+20AA) (see method for details) in M9Glc medium. Two biological replicates each with two technical replicates were performed and presented. **(e)** Normalized lag phase length of the *ΔrelAΔspoT207::cat* (ppGpp^0^) strain without or with supplemented arginine (5.2 mM, R), uracil (0.2 mM, U), glutamate (0.6 mM, E) or their varied combinations, relative to the *spoT^wt^* strain in M9Glc medium. Red dots indicate fails of regrowth within the tested time range of 72 hrs.

**Extended Data Fig. 4.**
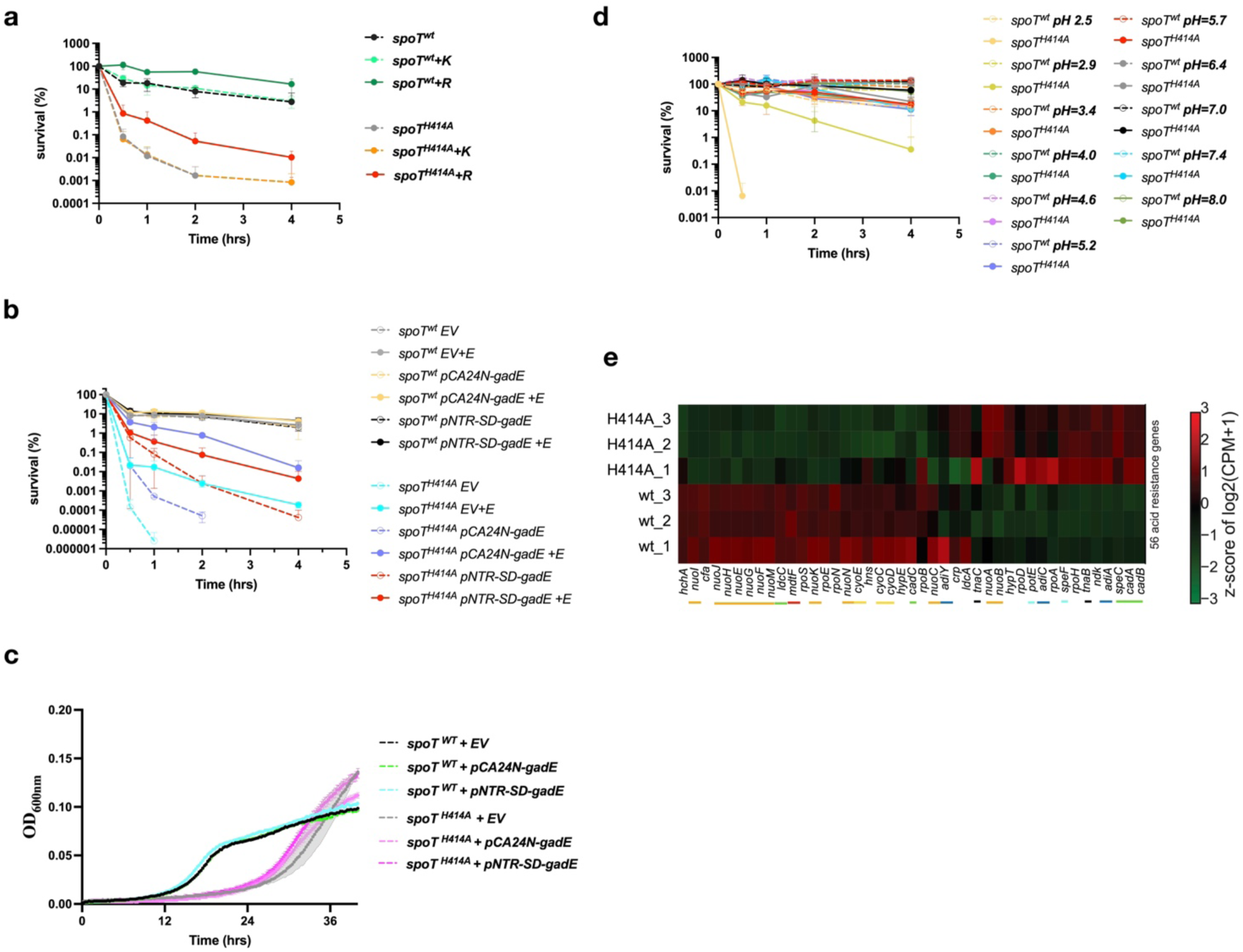
**(a)** Normalized percentage of surviving cells of various strains after challenged with pH=2.5. The data were normalized to the starting CFUs of each respective tested strain. +K, +R, lysine (0.6 mM), arginine (0.6 mM) included in the pH=2.5 medium, respectively. **(b)** Normalized percentage of surviving cells of *spoT^wt^* or *spoT^H414A^* strains containing various plasmid vectors (pCA24N ^42^, pNTR-SD ^43^) after challenged with pH=2.5. The data were normalized to the starting CFUs of each respective tested strain. EV, empty vector pCA24N. +E, glutamate (0.6 mM) included in the pH=2.5 medium. **(c)** Representative growth curves of *spoT^H414A^* and *spoT^wt^* strains containing either the empty vector (EV) or a *gadE* gene on pCA24N or pNTR-SD vectors in M9Glc medium. Two biological replicates each with two technical replicates were performed and presented. **(d)** Normalized percentage of surviving cells of *spoT^wt^* and *spoT^H414A^* strains after challenged with media of various pH values. The data were normalized to the starting CFUs of each respective tested strain. **(e)** Heatmap of genes involved in acid resistance. Colored bars indicate genes belonging to the same acid resistance systems: orange, *nuo* genes encoding NADH dehydrogenase I; yellow, *cyo* genes encoding cytochrome *bo* oxidase; green, lysine dependent acid resistance systems; cyan, ornithine dependent acid resistance system; black, alkaline stress; dark blue, arginine dependent acid resistance system.

**Extended Data Fig. 5.**
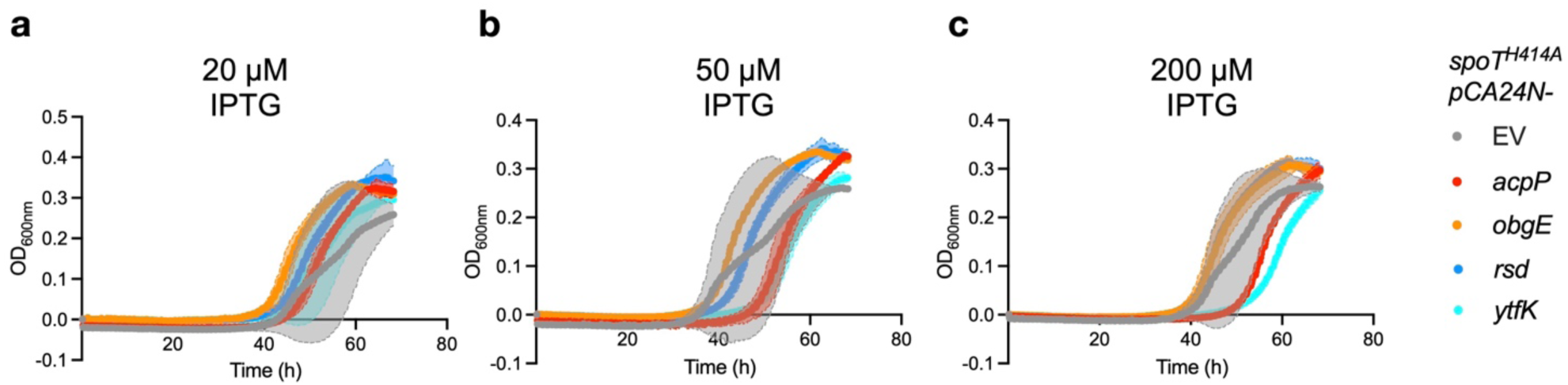
Growth dynamics of *spoT^H414A^* strains expressing candidate SpoT regulators. Growth curves of *spoT^H414A^* strains carrying either the empty vector (EV) pCA24N or pCA24N derivatives expressing *acpP, obgE, rsd,* or *ytfK*. Cells were grown in M9Glc medium supplemented with the indicated concentrations of IPTG to induce expression of the respective genes. Data represent the mean ± s.d. from three independent biological replicates.

**Extended Data Fig. 6.**
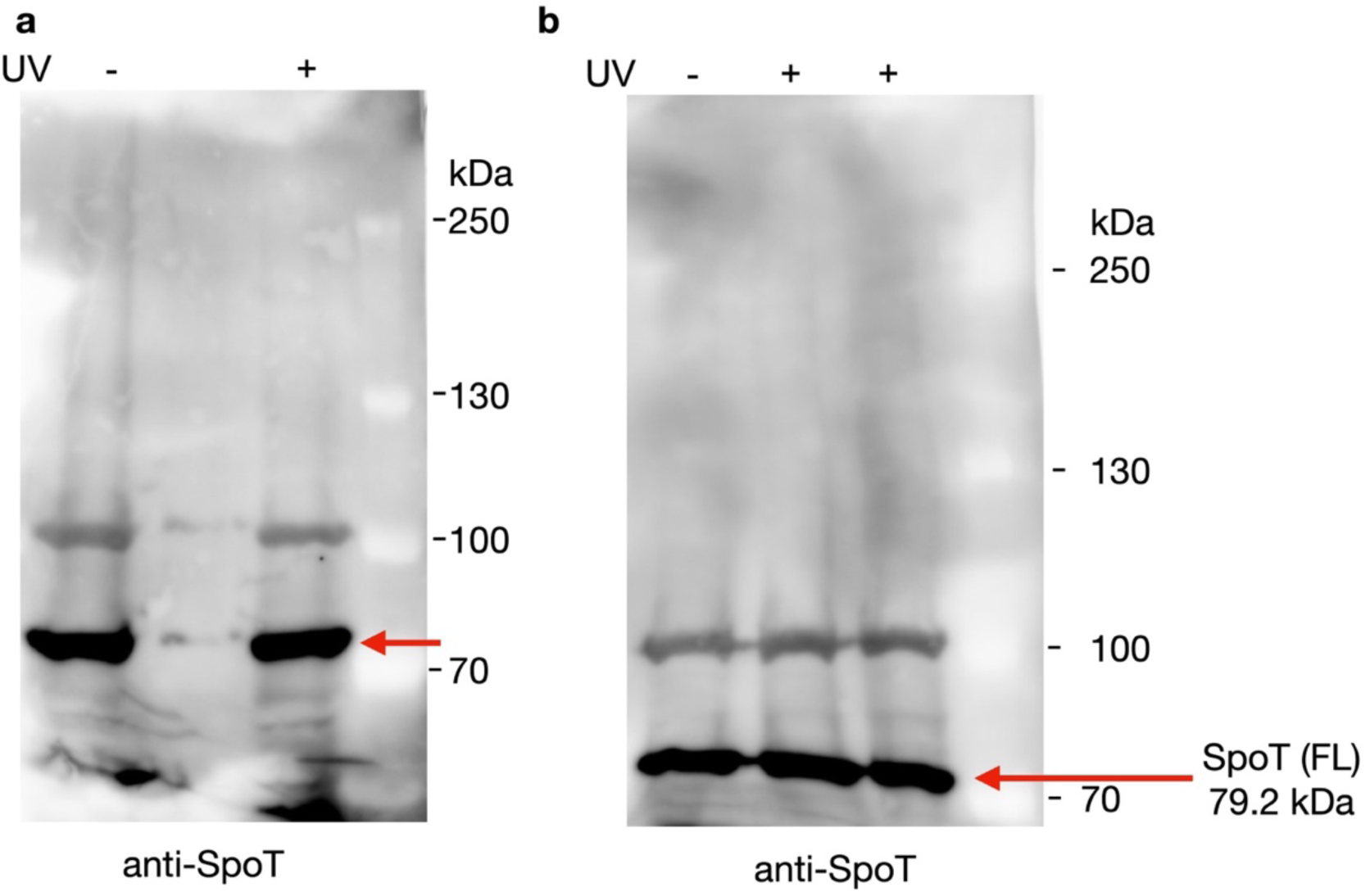
Western blot analysis of SpoT following unnatural amino acid-mediated UV crosslinking. SpoT carrying a photo-crosslinkable unnatural amino acid at position 414 was expressed in a *ΔrelA ΔspoT* background and subjected to UV irradiation (365 nm) to induce covalent crosslinking to proximal proteins. Samples collected in the absence (-) or presence (+) of UV irradiation were analyzed by SDS-PAGE and immunoblotting using anti-SpoT antiserum. Two independent representative experiments are shown (**a, b**). The positions of full-length SpoT are indicated by the red arrows.

**Extended Data Fig. 7.**
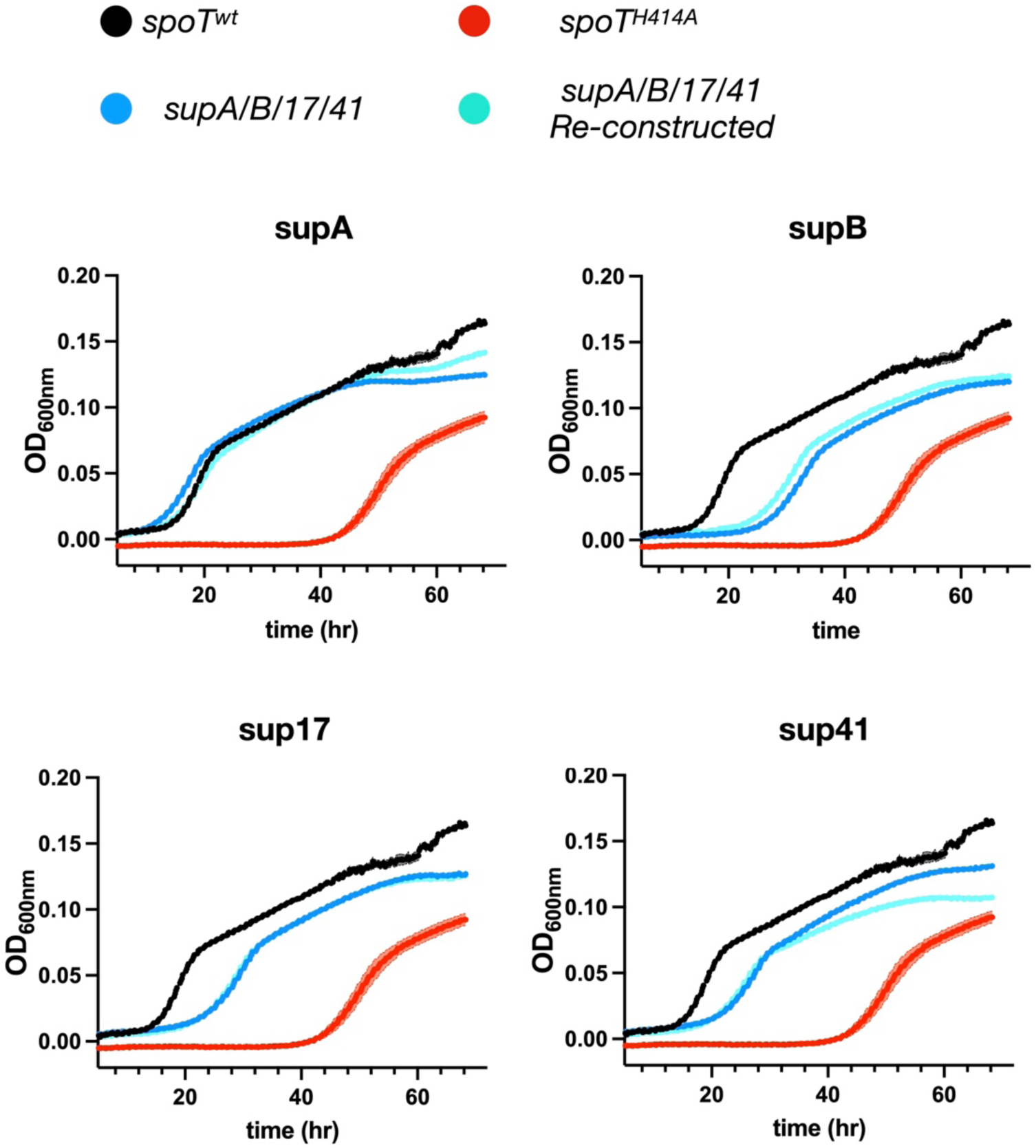
Growth dynamics of re-constructed four suppressors (cyan) (*supA/supB/sup17/sup41*) in comparison to *spoT^H414A^*, *spoT^wt^*, and the four random suppressors (blue).

**Extended Data Fig. 8.**
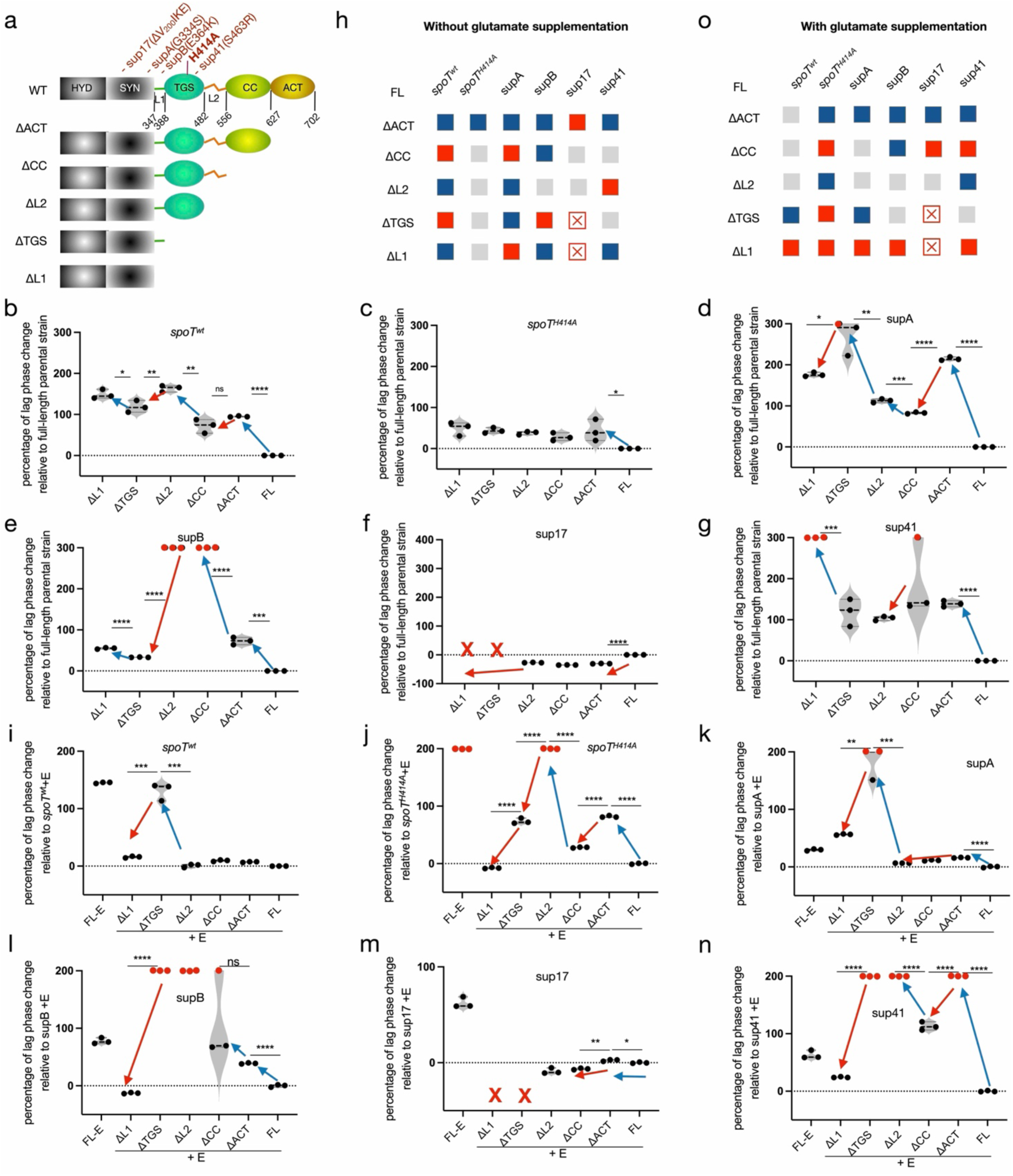
Distributed C-terminal domain regulation tunes SpoT-dependent growth under basal conditions. **(a)** Schematic representation of sequential truncations of SpoT C-terminal regulatory domains introduced into different genetic backgrounds. **(b-g)** Normalized lag phase duration during growth in M9Glc minimal medium, expressed as a percentage relative to the corresponding full-length (FL) SpoT parental strain. Red dots indicate strains that failed to resume growth within the 72-h observation period. Red crosses indicate truncation variants that could not be obtained. **(h)** Summary of the effects of individual domain deletions on growth resumption in M9Glc, based on data in panels **b-g**. Blue and red blocks indicate positive or negative effects on growth, respectively. Red crosses denote non-viable genotypes, and grey blocks indicate conditions in which no clear effect could be inferred due to indistinguishable growth behavior. **(i-n)** Normalized lag phase duration during growth in M9Glc supplemented with glutamate (0.6 mM), expressed relative to the corresponding FL parental strains. **(o)** Summary of domain-specific effects on growth resumption in the presence of glutamate, based on panels **i-n**. Color coding is as in panel **h**.

**Extended Data Fig. 9.**
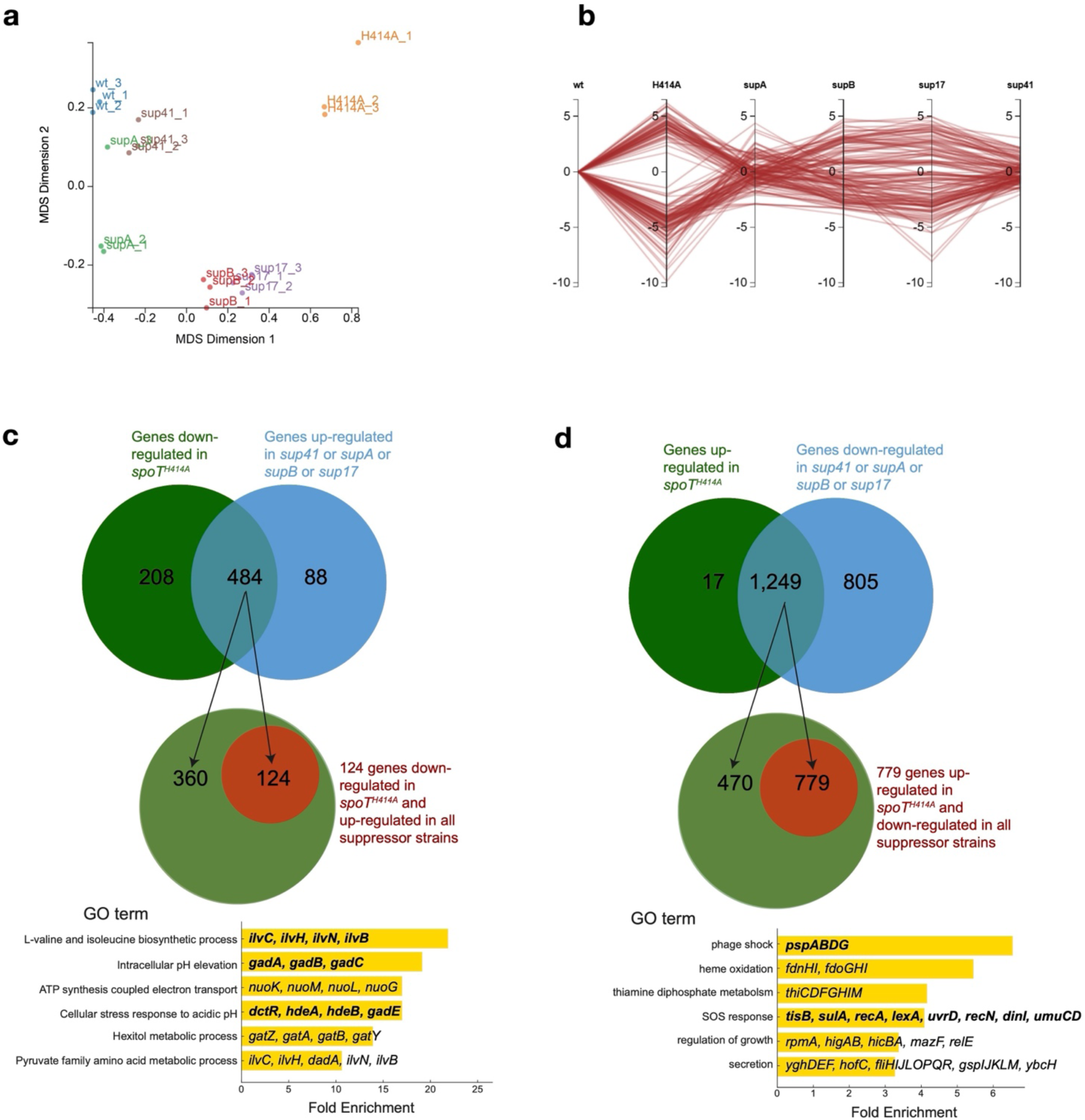
Suppressor mutations partially restore global transcriptional programs disrupted in *spoT^H414A^*cells. **(a)** Parallel coordinates plot showing genes with strong differential expression (|log₂ fold change| > 4, FDR < 0.05) across *spoT^wt^*, *spoT^H414A^* and four intragenic suppressor strains. (**b**) Multidimensional scaling (MDS) plot of transcriptomic profiles from the same six strains, with individual biological replicates shown. Growth conditions and sampling strategy are described in the Materials and Methods. Panels **a** and **b** were both generated by DEGUST. **(c)** Cross-suppressor analysis of genes downregulated in *spotT^H414A^*. Top, Venn diagram showing overlap of genes downregulated in *spotT^H414A^*and upregulated in one or more suppressor strains. Middle, subset of genes downregulated in *spotT^H414A^* and consistently upregulated in all four suppressor strains. Bottom, Gene Ontology (GO) enrichment analysis of the 124 genes restored across all suppressors. All genes are listed in **Table S4. (d)** Cross-suppressor analysis of genes upregulated in *spotT^H414A^*. Top, Venn diagram showing overlap of genes upregulated in *spotT^H414A^*and downregulated in one or more suppressor strains. Middle, subset of genes upregulated in *spotT^H414A^* and consistently downregulated in all four suppressor strains. Bottom, GO enrichment analysis of the 779 genes consistently repressed across all suppressors. Genes were extracted from DEGUST using thresholds of |log₂ fold change| > 1 and FDR < 0.05. Gene lists are provided in **Table S4.** Within GO term annotations, gene names shown in bold indicate transcripts with particularly strong differential expression.

**Extended Data Fig. 10.**
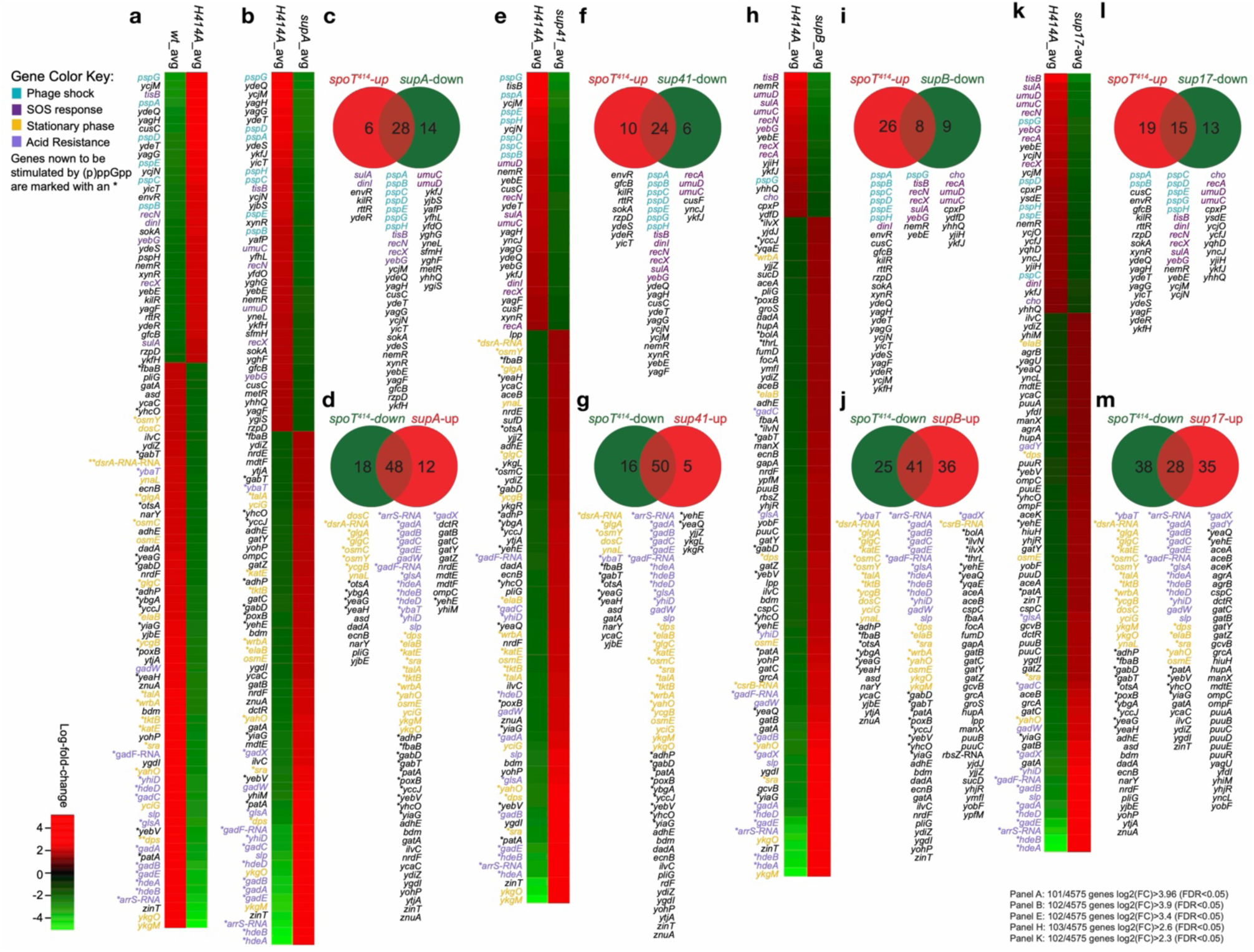
Suppressor mutations differentially correct transcriptional dysregulation in *spoT^H414A^* cells. **(a)** Heatmap of genes differentially expressed in *spoT^H414A^*relative to *spoT^wt^*. DEGUST analysis using thresholds of |log₂ fold change| > 3.96 and FDR < 0.05 identified 100 genes from a total of 4,575 genes analyzed. Highlighted gene classes include phage shock response genes (*psp*, blue), SOS response genes (magenta), acid stress response genes (deep purple), and stationary-phase-induced genes (gold). Genes marked with an asterisk (*) denote transcripts previously reported to be ppGpp-inducible. (**b, e, h, k)** Heatmaps comparing *spoT^H414A^*with each suppressor strain. FDR thresholds were identical to those used in panel (**a**), while |log₂ fold change| cutoffs were adjusted to return approximately 100 genes per comparison. DEGUST identified 102 genes for supA (|log₂ fold change| > 3.9), 97 genes for sup41 (|log₂ fold change| > 3.4), 94 genes for supB (|log₂ fold change| > 2.6), and 91 genes for sup17 (|log₂ fold change| > 2.3). Gene lists for each comparison are provided in **Table S5**. **(c, d, f, g, i, j, l, m)** Venn diagram analyses showing overlap of genes dysregulated in *spoT^H414A^*and in each suppressor strain. Genes differentially expressed in *spoT^H414A^*relative to *spoT^wt^* (from panel **a**) were used as the reference gene set for all Venn analyses, with upregulated and downregulated genes analyzed separately.

**Extended Data Fig. 11.**
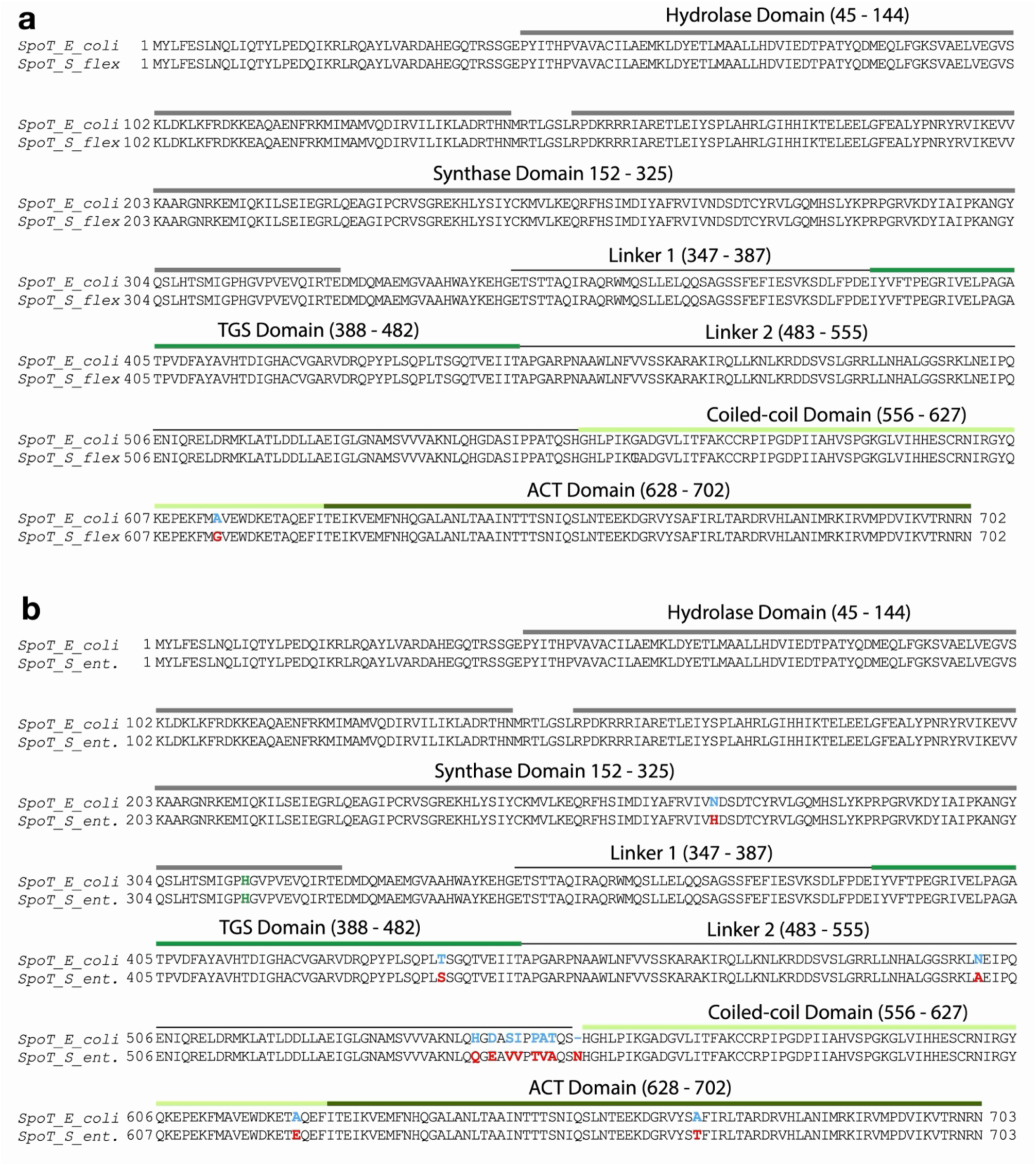
Sequence conservation and divergence of SpoT in enteric bacteria. Pairwise sequence comparison of *Escherichia coli* SpoT_ec_ with SpoT homologs from *Salmonella enterica* serovar Typhimurium SL1334 (SpoT_st_) and *Shigella flexneri* (SpoT_sf_). SpoT_sf_ is nearly identical to SpoT_ec_, differing by a single conserved substitution (A614G) within the CC domain. SpoT_st_ (702 amino acids) shares 98.15% overall sequence identity with SpoT_ec_ (701 amino acids), but contains several amino acid substitutions distributed across multiple domains, including the hydrolase (HYD; N267H), TGS (T437S), linker-2 (L2; nine substitutions: N501A, H542Q, D544E, S546V, I547V, P549T, A550V, T551A, and an insertion of N between residues 553-554), CC (A621E), and ACT (A668T) domains.

## ACKNOWLEDGEMENTS

We thank Paulina Grucella for helping with Western blot. The work is supported by a Novo Nordisk Foundation Project Grant (NNF19OC0058331), a Danmarks Frie Forskningsfond grant (2032-00030B) to Y.E.Z, the European Union’s Horizon 2020 research and innovation programme under the Marie Skłodowska-Curie grant agreement (N° 801199) to M.L.S, and the Chinese Scholarship Council program (N° 202206870009) to Y.Y.L.

## AUTHOR CONTRIBUTIONS

Y.E.Z designed this study; Y.Y.L, M.L.S and Y.E.Z acquired fundings; Y.Y.L, M.L.S, Y.E.Z acquired experimental data; X.C.Z, L.D, K.G, Y.E.Z performed the bioinformatics analyses; Y.E.Z, K.G, wrote the draft manuscript; and all authors analysed the data and edited the manuscript.

## EXPERIMENTAL MODEL AND SUBJECT DETAILS

*E. coli* DH5α (Stratagene) was used for cloning and *E. coli* K-12 MG1655 for experiments requiring an *E. coli* wt strain background (see **Table S1** for further details). For cloning, nutrient broth (Oxoid), with agar and antibiotics (chloramphenicol (25 µg/ml), ampicillin (100 µg/ml), and/or kanamycin (25 µg/ml)) when appropriate, was used. The different MG1655 strains were cultivated in Lysogeny Broth (LB) as well as M9Glc media (1x M9 salt, pH 7.4 1 mM MgSO4, 0.1 mM CaCl_2_, Thiamine (0.001 mg/1 ml), 0.2 % (g/ml) glucose) at 37 °C with agitation (160 rpm).

## METHOD DETAILS

### Isolation of suppressor mutations rescuing the growth of *spoT^H414A^* in M9Glc

The *spoT^H414A^* (YZ527) strain was grown overnight in LB media and then washed twice with PBS buffer. Approx. 10^7^ cells were plated on one M9Glc plate and incubated at 30°C. Colonies of different sizes were selected between 24-72 hours of incubation, and the restored growth was confirmed in solid and liquid M9Glc medium.

### Growth assay in liquid minimal media

To measure the growth curves of various strains in M9Glc media, the OD_600nm_ was measured of overnight cultures grown in LB of each strain, washed twice with 1x Phosphate Buffered Saline (PBS) and normalised in PBS and inoculated into fresh M9Glc media with a starting OD_600nm_ = 0.05 unless otherwise stated. The growth was measured every 15 min for 72 hrs in a plate reader (BioTek) presetted at 30°C with double orbital agitation at 548 cpm. For strains carrying ASKA plasmids (*acpP, rsd, ytfK,* or *obgE*), protein expression was induced by addition of IPTG at final concentrations of 20, 50, or 200 µM, as indicated.

### Anti-serum against SpoT C-terminal domains

Because full-length SpoT proved difficult to obtain in a soluble form, we instead generated a truncated SpoT construct encompassing the C-terminal regulatory domains. A DNA fragment corresponding to the *spoT* gene was first amplified using primers pYZ12 and pYZ13 and cloned into the NcoI and BamHI restriction sites of the pET28b expression vector. Subsequently, primers pYZ341 and pYZ342 were used for quick-change mutagenesis to delete the N-terminal catalytic domains, yielding a construct encoding the TGS, CC, and ACT domains of SpoT fused to an N-terminal His₆-TEV tag (pET28b-his.tev.spoT(TGS-CC-ACT)). The resulting plasmid was transformed into *Escherichia coli* BL21(DE3) cells for protein expression. His₆-TEV-SpoT(TGS-CC-ACT) was purified under denaturing conditions using nickel-affinity chromatography. Cell pellets were resuspended in lysis buffer containing 6 M urea (pH 8.0), and clarified lysates were applied to Ni-NTA resin. Bound protein was eluted with buffer containing 50 mM Tris-HCl (pH 8.0), 150 mM NaCl, 5% (v/v) glycerol, 0.5 M imidazole, and 6 M urea. Purified His₆-TEV-SpoT(TGS-CC-ACT) protein was used as antigen to generate polyclonal antiserum in rabbits by Covalab (France), according to standard immunization protocols.

### Unnatural amino acid**-**mediated UV crosslinking

#### Principle of unnatural amino acid-mediated UV crosslinking

Site-specific incorporation of a photo-crosslinkable unnatural amino acid was used to identify potential protein interaction partners of SpoT. A diazirine-containing lysine analog (DiZPK) was incorporated at position H414 of SpoT via amber stop codon suppression, enabling UV-induced covalent crosslinking to proximal proteins under native growth conditions.

#### Plasmid construction

To generate an amber codon substitution at position H414 of SpoT, the *spoT* gene was amplified from *Escherichia coli* MG1655 genomic DNA using primer pairs pYZ347/348 and pYZ349/350. Overlap PCR was performed to replace the codon encoding H414 with a TAG stop codon. The resulting PCR product was digested with BamHI and EcoRI and cloned into the corresponding sites of the pNDM220 expression vector, generating pNDM220-RBS-His₆.spoT(H414TAG).His₆. To remove the native N-terminal histidine tag sequence, primers pYZ700 and pYZ701 were used to amplify the N-terminal region of *spoT* from MG1655 genomic DNA. This fragment was digested with BamHI and PmlI and cloned into the BamHI/PmlI sites of the intermediate pNDM220-RBS-His₆.spoT(H414TAG).His₆ plasmid, yielding the final construct pNDM220-RBS-spoT(H414TAG).His₆.

### Cell growth and unnatural amino acid incorporation

The *E. coli ΔrelA ΔspoT* strain harboring pNDM220-RBS-spoT(H414TAG).His₆ and pSupAR-MbPylRS(DiZPK) (YZ1350) was grown overnight in LB medium and subcultured 1:50 into fresh LB supplemented with 1% (w/v) glucose. Cultures were grown at 30 °C with shaking until OD_600nm_ ≈ 0.2, at which point DiZPK (MedChemExpress; CAS 1337883-32-5) was added to a final concentration of 100 µM. Cells were incubated for an additional 30 min with shaking at 160 rpm, followed by induction of SpoT expression with 1 mM IPTG for 1 h. Cells were then harvested by centrifugation, washed twice with phosphate-buffered saline (PBS), and resuspended in M9Glc medium. Cultures were incubated for an additional 30 min to allow recovery under minimal medium conditions prior to UV crosslinking.

### UV crosslinking

Cell density was measured, and an aliquot (1 ml) was collected prior to UV exposure as a non-crosslinked control. Samples were pelleted by centrifugation (13,000 rpm, 2 min), supernatants were removed, and cell pellets were flash-frozen. For crosslinking, cultures were exposed to UV light at 365 nm for 30 min using an F8T5BL UV lamp. Following irradiation, cells were harvested by centrifugation, washed once with PBS, pelleted, and stored at −20 °C until analysis.

### Protein extraction and immunoblotting

Cell pellets were resuspended in Laemmli sample buffer and normalized to equivalent OD_600nm_ values. Proteins were separated by SDS-PAGE using a 5% resolving gel and transferred to PVDF membranes. Membranes were probed with anti-SpoT antiserum (1:2,000 dilution) for 1 h at room temperature, followed by incubation with HRP-conjugated anti-rabbit secondary antibodies. Immunoreactive bands were detected by chemiluminescence using Pierce® ECL Western Blotting Substrate (Thermo Scientific, #32209) and imaged with an ImageQuant LAS4000 system.

### Measurements of (p)ppGpp by autoradiography

(p)ppGpp is a highly labile molecule not amenable to HPLC or mass spectrometry analysis. Therefore, thin layer chromatography (TLC) was used to quantify both pppGpp and ppGpp. The excess phosphates in M9Glc affect efficient labelling of nucleotides by H_3_^32^PO_4_, therefore MOPS-M9Glc hybrid media was prepared wherein the PBS buffer in M9Glc was replaced by the MOPS buffer of the MOPS media^44^ (i.e., containing MOPS-tricine 40 mM pH 7.2, 1 mM MgSO4, 0.1 mM CaCl2, Thiamine (0.001 mg/1 ml), 0.2 % (g/ml) glucose, NH_4_Cl 0.001 g/ml, 8.5 mM NaCl, 0.2 mM KHPO_4_). The strains were grown for ca. 18 hrs in MOPS-M9Glc media at 37 °C with agitation. Afterwards, the strains were adjusted to the OD_600nm_ of 0.05 in MOPS-M9Glc and grown at 37 °C with agitation until OD_600nm_ of 0.2-0.3 was reached. Then, 100 μl cells were transferred to a 2-ml Eppendorf tube together with 5 μl of 0.005 mCi H_3_^32^PO_4_ (100 μCi/ml, PerkinElmer). After 1h at 37 °C with 600 rpm agitation, 90 μl were transferred to a new tube with 20 μl 2N formic acid, and the sample was stored at -20°C until analysed by TLC as described in^18^.

### Sample preparation for RNA-sequencing

The strains were grown overnight in LB media before washed twice in PBS and inoculated into M9Glc media (OD_600nm_ = 0.05). The strains were grown in M9Glc media until the early exponential phase (approximately OD_600nm_ = 0.2) before the cells were collected, and snap frozen with liquid nitrogen and stored at -80 °C. As the strain *spoT^H414A^* is unable to grow in M9Glc media, the starting OD_600nm_ was set to 0.2, and the cells were collected when the other strains are ready to harvest, i.e. ca. 18 hrs.

Total RNA was extracted using the RNeasy Mini Kit (Qiagen; 74104). Ribosomal RNA (rRNA) was depleted from purified total RNA with Ribo-Zero Plus rRNA Depletion kit (Illumina; 20040892). First and second strand complementary DNA (cDNA) were synthesized by cDNA Synthesis kit (Illumina; 20040895). RNA Index Anchors (Illumina; 20040899) was used for ligation of pre-index anchors to the ends of the double-stranded cDNA fragments to enable dual indexing. Barcoded cDNA libraries were constructed from anchor-ligated DNA fragments and sequenced on Illumina NovaSeq 6000 Sequencing System from Genewiz (Azenta Life Sciences), yielding a median depth of 39,619,482 paired-end 150 bp reads per sample.

### Strain construction

All strains and oligonucleotides used in this study are listed in **Tables S1** and **S2**, respectively. All chromosomal *in situ* mutations of *spoT* alleles were made by using the I-SceI-based λ -Red recombineering method^45^.

### The *spoT* E319Q and H72A D73A mutant strains

Briefly, the pWRG99 plasmid was transformed into the target strains (YZ38, YZ359, YZ543, YZ544, YZ545, YZ546) creating the target strains YZ313, YZ345, YZ826, YZ827, YZ828, and YZ829. The SceI site of the pWRG100 vector was PCR amplified using the primers pYZ611 and pYZ612 for the E319Q mutation and the primers pYZ617 and pYZ618 for the H72A D73A mutation. These PCR DNAs were then electroporated into the above strains (YZ38, YZ359, YZ543, YZ544, YZ545, YZ546). Then, the oligos pYZ613 and pYZ616 were used to introduce the E319Q and H72A D73A mutations into the specific *spoT* alleles *in situ*, generating the E319Q mutant strains (YZ872, YZ873, YZ874, YZ875, YZ876) and H72A D73A mutant strains (YZ900, YZ902, YZ904, YZ905).

### The C-terminal serial truncated mutant strains of *spoT*

pYZ796 and pYZ797 were used to amplify the SceI from pWRG100 and introduced into YZ313, YZ345, YZ826, YZ827, YZ828, and YZ829. The serial truncations were performed using the repairing oligos pYZ798, pYZ799, pYZ901, pYZ801, pYZ902, deleting the according domains (ACT, ACT-CC, ACT-CC-Linker2, ACT-CC-Linker2-TGS, ACT-CC-Linker2-TGS-Linker1, respectively).

### Reconstruction of the *spoT^H414A^* suppressor mutations

Primers pYZ619 and pYZ620 were used to amplify the SceI DNA from pWRG100 vector before it was introduced into the strain *spoT^wt^* (YZ313). Then, the primers pYZ328 and pYZ70 were used to amplify the corresponding *spoT* alleles using the chromosomal DNA of the strains supA (YZ543), *supB* (YZ544), sup17 (YZ545), and sup41 (YZ546) respectively as the templates. These DNAs were used to introduce the specific *spoT* suppressing mutations.

#### P1 phage transduction

The *spoT* genes of the suppressor strains YZ543, YZ544, YZ545, and YZ546 were inactivated via P1 phage transduction. The donor strain for making the P1 phage lysate was the *spoT207::cat* strain (YZ47)^13^, which contains a chloramphenicol resistance cassette together with *spoT207* deletion allele for selection. The phage lysate was used to infect strains YZ543, YZ544, YZ545, and YZ546 and the chloramphenicol resistant transductants were selected and confirmed via diagnostic PCR.

### Plasmids constructions

To construct *E. coli (Ec), Salmonella typhimurium(st) or Shigella flexneri(sf) spoT* wild type (wt) and H414A mutant plasmids, standard molecular cloning procedures were used. For *E. coli*, the *spoT* coding sequence was PCR-amplified from MG1655 *spoT^wt^* (YZ38) or *spoT^H414A^*(YZ527) using primers pYZ1201 and pYZ1202. The pBAD33 vector was isolated from *E. coli* DH5α (YZ593). Both the amplified *spoT* fragments and pBAD33 were digested with *KpnI* and *HindIII* and ligated using T4 DNA ligase to generate pBAD33- *spoT^wt^* and pBAD33- *spoT^H414A^*. For *S. Typhimurium*, the H414A mutation was introduced into pGEN-MCS-*spoT_St_* (YZ692)^46^ by site-directed mutagenesis using primers pYZ1312 and pYZ1313, yielding pGEN-MCS-spoT_st_ (H414A). For *S. flexneri*, an A614G substitution was introduced by overlap-extension PCR using pBAD33-*spoT_Ec_*(wt) or pBAD33- *spoT_Ec_*(H414A) as templates. Upstream and downstream fragments were amplified with primer pairs pYZ1201/pYZ1346 and pYZ1202/pYZ1345, respectively, fused by overlap PCR using primers pYZ1201 and pYZ1202, and cloned into pBAD33 following digestion with *KpnI* and *HindIII* to generate pBAD33-*spoT*_Sf_(WT) and pBAD33-*spoT_Sf_*(H414A). To construct the plasmid pCA24N-argA(H15Y), quick-change mutagenesis was carried out using pCA24N-argA(wt) as template and the primers pYZ1168 and pYZ1169. All plasmids were transformed into *E. coli* DH5α (see **Table S1**), purified using a commercial miniprep kit (NEB-T1110L), and confirmed by sequencing with primers pYZ404, pYZ405, and pYZ71 (for pBAD33 constructs), pYZ1305, pYZ1306, and pYZ1228 (for pGEN-MCS constructs) or pYZ34 and pYZ35 (for pCA24N constructs; Eurofins Genomics).

### Acid resistance assay

Overnight (16 hrs) cultures were grown in LB medium at 37°C with shaking (160 rpm). Cells were harvested by centrifugation, washed twice with PBS, and normalized to an optical density at 600 nm (OD_600nm_) of 0.4 in minimal medium supplemented with 0.2% glucose (M9Glc, pH 7.4). Cultures were incubated at 37°C with shaking (160 rpm) for 30 min to allow adaptation to M9Glc. After adaptation, 1 mL cells were pelleted and resuspended in the same volume of M9Glc medium pre-adjusted to various pH values (from 2.5 to 8.0), with or without supplemented 0.6 mM glutamate (or other amino acids). Acid challenge was then performed and assessed at 37°C with shaking (160 rpm). Cell viability was determined at time zero (i.e., before shifting to M9Glc with a low pH), and at 0.5, 1, 2 and 4 h after acid challenges. For viable-cell counts, 40 µL cell culture at each time point was collected and 10-fold serially diluted (10⁰-10⁻⁷) in LB medium, and 5 µL of each dilution was spotted on LB agar plates, allowing colony formation and enumeration and calculation of CFUs the next day.

### Western blot

Western blot analysis of protein levels of various mutant SpoT was performed as reported^18^. Briefly, cells of various strains were collected the same manner as preparation for RNA sequencing (see above, and **Fig. S1C**). These cells were normalised to have an equal OD_600nm_ per ml and mixed with 2x SDS-PAGE loading buffer. Samples were heated for 15 min at 95°C and 20 ul of each were separated in 10% SDS-PAGE gels. The proteins were transferred to a PVDF membrane (Amersham) and the SpoT proteins detected by first incubation with anti-SpoT antiserum (Covalab) and then the secondary HRP conjugated rabbit IgG antibody (SIGMA). The SpoT bands were visualized by using the Pierce ECL chemiluminescence substrate (Thermo) and the signals were detected by an Imagequant LAS4100.

### Bioinformatic analysis

#### Preparing RNA-seq data

RNA-seq data (fastq files) were mapped by the BWA-MEM tool of Galaxy^47^ and then analysed by the featureCounts tool in Galaxy https://usegalaxy.eu/). The tabular format of the featureCounts files were exported from Galaxy and assembled into one single tabular file from which sub-files containing collections of genes of interest, e.g. amino acid biosynthesis, purine de novo synthesis, were extracted.

#### Analysis of RNA-seq data

To generate heatmaps and fold-of-changes, the tabular files containing the RNA-seq counts were analysed by DEGUST (https://degust.erc.monash.edu/). To generate Venn diagrams, gene lists were analysed by the jVenn application at (https://jvenn.toulouse.inrae.fr/app/index.html). Gene Ontology terms were generated by AmiGO (https://amigo.geneontology.org/amigo) and Panther (https://pantherdb.org/webservices/go/overrep.jsp). The json-files generated by PANTHER were analysed by ChatGPT that generated editable SVG files.

#### Other Bioinformatic Procedures

Protein sequence alignment was accomplished using JalView^48^, protein tertiary structure modelling and visualization by AlphaFold 3 and ChimeraX, respectively^49,50^.

## QUANTIFICATION AND STATISTICAL ANALYSIS

Signals of (p)ppGpp from autoradiography were quantified using ImageQuant (GE Healthcare). At least three biological replicates were performed and one representative shown in the **Figure**s.

For other physiological growth experiments, at least two biological replicates each with three technical replicates were done (see **Figure** legends for details). The length (hours) of lag phases was defined as the period from the growth start to the time point where the slope of growth curves first reaches the exponential-phase growth rate. These lag phage length values were then normalized to the parental strains or conditions, and plotted (see each of the legends for details). The student *t*-test was used for statistic analysis wherever applicable.

## DATA AVAILABILITY

The RNA-seq raw data produced in this study are available through the NCBI Sequence Read Archive under BioProject accession number PRJNA1097790.

## Legends to Supplementary Tables

**Table S1. Bacterial strains used in this study**

**Table S2. Oligonucleotides used in this study**

**Table S3. Differential gene expression analysis of RNA-seq data from *spoT^wt^* and *spoT^H414A^* strains .** In total, the transcription levels of 4575 genes of MG1655 were analysed. At a setting of |Log2(FC)| > 1 and FDR < 0.05, 865 and 949 genes were down and up-regulated, respectively. The change-of-folds |Log2(FC)| of each gene are also shown.

**Table S4. Gene ontology analysis of the four suppressor strains.**

**Sheet #1 (related to Extended Data Fig. 7C):** Genes downregulated in the *spoT^H414A^* strain relative to the *spoT^wt^* strain (Column A). Genes upregulated in either of the four suppressor strains *supA*, *sup41*, *supB* and *sup17* relative to the *spoT^H414A^* strain (Columns B to E). The list of genes in Columns B to E were used to generate a list of genes that were upregulated in at least one suppressor strain (Column F) and a list of genes that were upregulated in all 4 suppressor strains (Column G).

**Sheet #2 (related to Extended Data Fig. 7D):** Genes upregulated in the *spoT^H414A^*strain relative to the *spoT^wt^* strain (Column A). Genes downregulated in either of the four suppressor strains relative to the *spoT^H414A^*strain. The list of genes in Columns B to E were used to generate a list of genes that were upregulated in at least one suppressor strain (Column F) and a list of genes that were downregulated in all 4 suppressor strains (Column G). Gene lists in **Table S4** were generated by DEGUST at a setting of |Log2(FC)| > 1 and FRD < 0.05.

**Table S5. Comparison of *spoT^H414A^* with suppressor strains (related to Extended Data Fig. 8).** The Table yields the folds-of-changes in the suppressor strains relative to *spoT^H414A^*. The Log2(FC) setting in DEGUST was chosen to yield around 100 genes up and downregulated in each heatmap Panels a, b, e, h and k of **Extended Data Fig. 8**.

