## Supplementary material for "Basal ppGpp signalling by SpoT integrates metabolism with acid resistance": Table S1

**Table S1: Bacterial strains used in this study**

| **Strain** | **Relevant features** | **Reference** |
| --- | --- | --- |
| YZ38 | MG1655 *ΔrelA* | laboratory stock |
| YZ47 | MG1655 *ΔrelA*251::*kan* *ΔspoT207*::*cat* | ^1^ |
| YZ313 | MG1655 *ΔrelA* pWRG99 | This study |
| YZ345 | MG1655 *ΔrelA* pWRG99; *spoT* codon H414A | This study |
| YZ359 | MG1655 *ΔrelA spoT* codon H414A | This study |
| YZ527 | MG1655 *ΔrelA spoT* codon H414A | This study |
| YZ543 | MG1655 *ΔrelA spoT* codon H414A, supA | This study |
| YZ544 | MG1655 *ΔrelA spoT* codon H414A, supB | This study |
| YZ545 | MG1655 *ΔrelA spoT* codon H414A, sup17 | This study |
| YZ546 | MG1655 *ΔrelA spoT* codon H414A, sup41 | This study |
| YZ554 | MG1655 *ΔrelA spoT* codon H414A, supA, *ΔspoT207::cat* | This study |
| YZ555 | MG1655 *ΔrelA spoT* codon H414A, supB, *ΔspoT207::cat* | This study |
| YZ556 | MG1655 *ΔrelA spoT* codon H414A, sup17, *ΔspoT207::cat* | This study |
| YZ557 | MG1655 *ΔrelA spoT* codon H414A, sup41, *ΔspoT207::cat* | This study |
| YZ593 | DH5α pBAD33 | laboratory stock |
| YZ692 | Salmonella typhimurium SL1344 ΔrelA ΔspoT pGEN-MCS-spoTst | ^2^ |
| YZ826 | MG1655 *ΔrelA* pWRG99; *spoT* codon H414A, supA | This study |
| YZ827 | MG1655 *ΔrelA* pWRG99; *spoT* codon H414A, supB | This study |
| YZ828 | MG1655 *ΔrelA* pWRG99; *spoT* codon H414A, sup17 | This study |
| YZ829 | MG1655 *ΔrelA* pWRG99; *spoT* codon H414A, sup41 | This study |
| YZ872 | MG1655 *ΔrelA* *spoT* codon H414A, supA, E319Q | This study |
| YZ873 | MG1655 *ΔrelA spoT* codon H414A, supB, E319Q | This study |
| YZ874 | MG1655 *ΔrelA spoT* codon H414A, sup17, E319Q | This study |
| YZ875 | MG1655 *ΔrelA spoT* codon H414A, sup41, E319Q | This study |
| YZ876 | MG1655 *ΔrelA spoT* codon H414A, E319Q | This study |
| YZ886 | MG1655 *ΔrelA* SceI::Cam replacing *spoT* excl. Start and stop codons | This study |
| YZ900 | MG1655 *ΔrelA spoT* codon H414A, supB, H72A D73A | This study |
| YZ902 | MG1655 *ΔrelA spoT* codon H414A, sup41, H72A D73A | This study |
| YZ904 | MG1655 *ΔrelA spoT* codon H414A, H72A D73A | This study |
| YZ905 | MG1655 *ΔrelA* H72A D73A | This study |
| YZ1227 | MG1655 *ΔrelA* *sceI::cam* integrated directly after TAA (of *spoT*) with an extra AA at the end of *sceI* cassette | This study |
| YZ1228 | MG1655 *ΔrelA* *sceI::cam* integrated directly after TAA (of *spoT*) with an extra AA at the end of *sceI* cassette, H414A | This study |
| YZ1229 | MG1655 *ΔrelA* *sceI::cam* integrated directly after TAA (of spoT) with an extra AA at the end of *sceI* cassette , supA | This study |
| YZ1230 | MG1655 *ΔrelA* *sceI::cam* integrated directly after TAA (of *spoT*) with an extra AA at the end of *sceI* cassette, supB | This study |
| YZ1231 | MG1655 *ΔrelA* *sceI::cam* integrated directly after TAA (of *spoT*) with an extra AA at the end of *sceI* cassette, sup17 | This study |
| YZ1232 | MG1655 *ΔrelA* *sceI::cam* integrated directly after TAA (of *spoT*) with an extra AA at the end of *sceI* cassette , sup41 | This study |
| YZ1268 | MG1655 *ΔrelA spoT^wt^* ACT deleted | This study |
| YZ1269 | MG1655 *ΔrelA spoT^wt^* ACT + CC deleted | This study |
| YZ1448 | MG1655 *ΔrelA spoT^wt^* ACT + CC + L2 deleted | This study |
| YZ1271 | MG1655 *ΔrelA spoT^wt^* ACT + CC + linker2+ TGS deleted | This study |
| YZ1449 | MG1655 *ΔrelA spoT^wt^* ACT + CC + L2+ TGS+ L1 deleted | This study |
| YZ1273 | MG1655 *ΔrelA* *spoT^H414A^* ACT deleted | This study |
| YZ1274 | MG1655 *ΔrelA* *spoT^H414A^* ACT + CC deleted | This study |
| YZ1450 | MG1655 *ΔrelA spoT^H414A^* ACT + CC + L2 deleted | This study |
| YZ1276 | MG1655 *ΔrelA* *spoT^H414A^* ACT + CC + linker2+ TGS deleted | This study |
| YZ1461 | MG1655 *ΔrelA spoT^H414A^* ACT + CC + L2+ TGS+ L1 deleted | This study |
| YZ1278 | sup41 ACT deleted | This study |
| YZ1279 | sup41 ACT + CC deleted | This study |
| YZ1458 | sup41 ACT + CC + L2 deleted | This study |
| YZ1281 | sup41 ACT + CC + linker2+ TGS deleted | This study |
| YZ1459 | sup41 ACT + CC + L2+ TGS+ L1 deleted | This study |
| YZ1283 | sup17 ACT deleted | This study |
| YZ1284 | sup17 ACT + CC deleted | This study |
| YZ1456 | sup17 ACT + CC + L2 deleted | This study |
| YZ1286 | supA ACT deleted | This study |
| YZ1287 | supA ACT + CC deleted | This study |
| YZ1464 | supA ACT + CC + L2 deleted | This study |
| YZ1400 | supA ACT + CC + L2+ TGS deleted | This study |
| YZ1471 | supA ACT + CC + L2+ TGS+ L1 deleted | This study |
| YZ1288 | supB ACT deleted | This study |
| YZ1289 | supB ACT + CC deleted | This study |
| YZ1454 | supB ACT + CC + L2 deleted | This study |
| YZ1291 | supB ACT + CC + linker2+ TGS deleted | This study |
| YZ1455 | supB ACT + CC + L2+ TGS+ L1 deleted | This study |
| YZ1584 | MG1655 *ΔrelA spoT::IsceI* excpet the TTG start and TAAsstop codons, pWRG99 | This study |
| YZ1585 | MG1655 *ΔrelA spoT* H414A supA (reconstituted strain) | This study |
| YZ1586 | MG1655 *ΔrelA spoT* H414A supB (reconstituted strain) | This study |
| YZ1587 | MG1655 *ΔrelA spoT* H414A sup17 (reconstituted strain) | This study |
| YZ1588 | MG1655 *ΔrelA spoT* H414A sup41 (reconstituted strain) | This study |
| YZ1754 | ASKA pCA24N-argA^wt^ | ^3^ |
| YZ1755 | DH5α pCA24N-argA^H15Y^ | This study |
| YZ1772 | DH5α pBAD33-spoT_Ec_ | This study |
| YZ1776 | DH5α pBAD33-spoT_Ec_^H414A^ | This study |
| YZ1876 | DH5α pGEN-MCS-spoT_st_^H414A^ | This study |
| YZ1942 | DH5α pBAD33-spoT_sf_ | This study |
| YZ1943 | DH5α pBAD33-spoT_sf_^H414A^ | This study |

1.	Xiao, H., Kalman, M., Ikehara, K., Zemel, S., Glaser, G., and Cashel, M. (1991). Residual guanosine 3',5'-bispyrophosphate synthetic activity of relA null mutants can be eliminated by spoT null mutations. J Biol Chem *266*, 5980-5990.

2.	Chau, N.Y.E., Perez-Morales, D., Elhenawy, W., Bustamante, V.H., Zhang, Y.E., and Coombes, B.K. (2021). (p)ppGpp-Dependent Regulation of the Nucleotide Hydrolase PpnN Confers Complement Resistance in Salmonella enterica Serovar Typhimurium. Infect Immun *89*. 10.1128/IAI.00639-20.

3.	Kitagawa, M., Ara, T., Arifuzzaman, M., Ioka-Nakamichi, T., Inamoto, E., Toyonaga, H., and Mori, H. (2006). Complete set of ORF clones of Escherichia coli ASKA library (A Complete Set of E. coli K-12 ORF Archive): Unique Resources for Biological Research. DNA Research *12*, 291-299. 10.1093/dnares/dsi012.
