## Supplementary material for "Basal ppGpp signalling by SpoT integrates metabolism with acid resistance": Table S2

**Table S2: Primers used in this study**

| **Number** | **Name** | **Sequence 5’ to 3’** |
| --- | --- | --- |
| pYZ7 | spoT-seq-R | CCGACCTTCAAGGGTCAACC |
| pYZ9 | spoT-seq-F | GTCAGTGGTCGCGAGAAGCATC |
| pYZ10 | T7-seq-F | taatacgactcactataggg |
| pYZ18 | spoT-seq-F | CGTTGAAATCATTACCGCTC |
| pYZ34 | pCA24Nf | GGCCCTTTCGTCTTCACCTC |
| pYZ35 | pCA24Nr | GGCAACCGAGCGTTCTGAAC |
| pYZ69 | ΔspoT-conf-F | gtcgtcgttaatcacaaagc |
| pYZ70 | ΔspoT-conf-R | catgcagacggtcagatcag |
| pYZ71 | spoT-seq-R | gcactgcataagcgaagtcg |
| pYZ328 | ΔspoT-conf-F | GCCAGGAACAGCAAGAGCAG |
| pYZ404 | pBAD33-seq-F | ACACTTTGCTATGCCATAGC |
| pYZ405 | pBAD33-seq-R | CAGACCGCTTCTGCGTTCTG |
| pYZ417 | spoT-seq-R | TGAACTTCATGTACGCCAAC |
| pYZ611 | 60mer-spoT-E319Q-F | tttgcacacctcgatgatcggcccgcacGGTgtgccggttCTAGACTATATTACCCTGTTATCCC |
| pYZ612 | 60mer-spoT-E319Q-F | tccgccatctggtccatatcttcggtacggatctggacctATTTAAATGGCGCGCCTTAC |
| pYZ613 | 80mer-spoT-E319Q-ISceI | tttgcacacctcgatgatcggcccgcacGGTgtgccggttcaggtccagatccgtaccgaagatatggaccagatggcgga |
| pYZ616 | 80mer-spoT-H72D73AA-ISceI | gatgaaactcgactatgaaacgctgatggcggcgctgctgGCTGCAgtgattgaagatactcccgccacctaccaggatatggaac |
| pYZ617 | 60mer-spoT-H72D73AA-invF | gatgaaactcgactatgaaacgctgatggcggcgctgctgATTTAAATGGCGCGCCTTAC |
| pYZ618 | 60mer-spoT-H72D73AA-invR | gttccatatcctggtaggtggcgggagtatcttcaatcacCTAGACTATATTACCCTGTTATCCC |
| pYZ619 | 60mer-spoT-clean del-F | tgctgaaggtcgtcgtTAAtcacaaagcgggtcgcccTTGCTAGACTATATTACCCTGTTATCCC |
| pYZ620 | 60mer-spoT-clean del-R | CGCagatgcgtgcataacgtgttgggttCATaaaacaTTAATTTAAATGGCGCGCCTTAC |
| pYZ701 | spoT-seq-R | CCGCTTGAACGTGTTTGC |
| pYZ702 | spoT-cloning-F1 | CAGGCGTATCTCGTTGCAC |
| pYZ703 | spoT-cloning-R1 | CACTGCATAAGCGAAGTCG |
| pYZ704 | spoT-cloning-F2 | CAT TGT CGA GCT GCC TG |
| pYZ705 | spoT-cloning-R2 | TCTTCCGTATTCAAACTTTGAATATTC |
| pYZ796 | 60mer-spoTTAAins-F | GATGCCAGACGTGATTAAAGTCACCCGAAACCGAAATTAACTAGACTATATTACCCTGTTATCCC |
| pYZ797 | 60mer-spoTTAAins-R | TTTCGCagatgcgtgcataacgtgttgggttCATaaaacaTTATTTAAATGGCGCGCCTTAC |
| pYZ798 | 80mer-spoTΔACT | tgtggaatgggataaagagacggcgcaggagTTCATCACCTAAtgttttATGaacccaacacgttatgcacgcatctGCG |
| pYZ799 | 80mer-spoTΔACTΔCC | TGGGGACGCCTCCATTCCACCGGCAACCCAAagccacggaTAAtgttttATGaacccaacacgttatgcacgcatctGCG |
| pYZ801 | 80mer-spoTΔ (ACL2) ΔTGS | tatcgagagcgttaaatccGATCTCTTCCCGgatgagattTAAtgttttATGaacccaacacgttatgcacgcatctGCG |
| pYZ901 | 80mer-spoTΔACTΔLinker2 | aattcgtcagttgctgaaaaacctcaagcgtGATGATTCTTAAtgttttATGaacccaacacgttatgcacgcatctGCG |
| pYZ902 | 81mer-spoTΔ (ACL2T) ΔLinker1 | gtgttgccgcgcactgggcttataaagagcacggcgaaaccTAAtgttttATGaacccaacacgttatgcacgcatctGCG |
| pYZ1168 | QCargAH15Y-R | ACCGAataGCGGAATCCCTCGAC |
| pYZ1169 | QCargAH15Y-F | ATTCCGCtATTCGGTTCCCTATATCAATACC |
| pYZ1201 | KpnI-spoT-F | ggGGTACCaTGTATCTGTTTGAAAGCCTGA |
| pYZ1202 | HindIII-EcoRI-spoT-R | cccAAGCTTgaattcTTAATTTCGGTTTCGGGT |
| pYZ1228 | pGEN-seq-F | aatcgtttgcactgtctctg |
| pYZ1305 | SL1344-spoT-seq-F | GCACCGTTTAGGTATTCATCAC |
| pYZ1306 | pGEN-seq-R | CAGATTTCGTGATGCTTGTC |
| pYZ1312 | QC-SL1344-spoT-H414A-F | ATGCAGTGgcTACCGACATCGGCCAC |
| pYZ1313 | QC-SL1344-spoT-H414A-R | TGTCGGTAgcCACTGCATAGGCAAAATCC |
| pYZ1345 | QCspoTecA614G-F | tttatggGTGTGGAATGGGATAAAGAGAC |
| pYZ1346 | QCspoTecA614G-R | ttccacaCCCATAAACTTCTCTGGCTCTTT |
